# BK polyomavirus evades innate immune sensing by disrupting the mitochondrial network and membrane potential and by promoting mitophagy

**DOI:** 10.1101/2020.03.19.994988

**Authors:** Julia Manzetti, Fabian H. Weissbach, Gunhild Unterstab, Marion Wernli, Helmut Hopfer, Cinthia B. Drachenberg, Christine Hanssen Rinaldo, Hans H. Hirsch

## Abstract

Immune escape contributes to viral persistence, yet little is known about human polyomaviruses. BK-polyomavirus (BKPyV) asymptomatically infects 90% of the human population, but causes early allograft failure in 10% of kidney transplants. Despite inducing potent virus-specific T-cells and neutralizing antibodies, BKPyV persists in the kidneys and regularly escapes from immune control as indicated by urinary shedding in immunocompetent individuals. Here, we report that BKPyV disrupts the mitochondrial network and its membrane potential when expressing the 66aa-long agnoprotein during late replication. Agnoprotein impairs nuclear IRF3-translocation, interferon-*β* expression, and promotes p62-mitophagy in vitro and in kidney transplant biopsies. Agnoprotein-mutant viruses unable to disrupt mitochondria show reduced replication, which can be rescued by type-I-interferon-blockade, TBK1-inhibition, or CoCl_2_ treatment. Agnoprotein is necessary and sufficient, using its amino-terminal and central domain for mitochondrial targeting and disruption, respectively. JCPyV- and SV40-infection similarly disrupt the mitochondrial network indicating a conserved mechanism facilitating polyomavirus persistence and post-transplant disease.

## Introduction

Viruses infect all forms of life, and while metagenomics are revealing an increasing complexity of viromes (Simmonds et al., 2017; Virgin, 2014), the underlying principle remains the same: viral genetic information present as DNA or RNA is decoded by host cells, thereby enabling a programmed take-over of cell metabolism to accomplish the essential steps of viral genome replication, packaging, and progeny release to infect new susceptible cells and hosts (Lynn W. Enquist, 2013). With progeny rates of ten- to one hundred-thousand per cell, virus replication represents a severe burden compromising host cell function and viability, and ultimately organ and host integrity. Host defense mechanisms include antiviral restriction factors (Kluge et al., 2015) as well as innate and adaptive immune responses (McNab et al., 2015) intercepting early and late steps of the viral life cycle. Given the obligatory intracellular step of virus replication, the innate immune response is faced with the challenge of discriminating virus and its regulatory and structural units as “non-self” amidst physiological host cell constituents. This task is partly accomplished by identifying pathogen-associated molecular patterns (PAMPs) such as repetitive protein, lipid, and sugar structures through pattern recognition receptors (PRRs). In the last decade, cytoplasmic sensing of nucleic acids has emerged as a key mechanism of intracellular innate immune sensing (Takeuchi and Akira, 2010). Evolutionarily linked to detecting DNA damage and the failing integrity of the nucleus or mitochondria, which may result from metabolic stress, toxicity, or cancer (Hartlova et al., 2015; Li and Chen, 2018; West and Shadel, 2017), viral RNA and DNA were found to be similarly sensed through PRRs such as RIG-I/MDA-5/MAVS and cGAS/STING (Goubau et al., 2013; Hartlova et al., 2015; McFadden et al., 2017). MAVS and STING have been located on the membranous platforms of mitochondria and the endoplasmic reticulum (ER). Besides direct cross-talk (Zevini et al., 2017) and proximity via mitochondria-associated ER-membranes, both pathways converge in inducing type-1 interferon expression following the activation of the downstream kinases TBK-1 and IKK*_ε_*, and the phosphorylation and nuclear translocation of transcription factors such as IRF3, IRF7, and NFkB (Liu et al., 2015; McFadden et al., 2017). Interferons are key mediators of the antiviral state in infected and neighbouring cells and help to shape activation of the adaptive antigen-specific immune responses (Iwasaki and Medzhitov, 2010; Schneider et al., 2014). Different molecular mechanisms of innate immune activation are induced in acute and chronic infections with human RNA or DNA viruses, and a variety of strategies has been described which permit transient or persistent viral evasion (Garcia-Sastre, 2017). However, such aspects are incompletely understood for small non-enveloped DNA viruses such as human polyomaviruses.

BK polyomavirus (BKPyV) is one of at least 10 human polyomaviruses and infects >90% of the general population typically during childhood without specific illness (Greenlee and Hirsch, 2017; James A. DeCaprio, 2013). Although BKPyV induces potent virus-specific CD4 and CD8 T-cells (Binggeli et al., 2007; Chen et al., 2008; Cioni et al., 2016; Leboeuf et al., 2017) and neutralizing antibodies (Pastrana et al., 2012; Shah et al., 1980; Solis et al., 2018), the virus latently persists in the kidneys and regularly escapes from immune control as indicated by asymptomatic urinary virus shedding in immunocompetent healthy individuals (Egli et al., 2009; Imperiale and Jiang, 2016; Zhong et al., 2007). In immunosuppressed patients, BKPyV-replication increases in rate and magnitude, progressing to nephropathy and haemorrhagic cystitis in 1%-15% and 5%-25% of kidney transplant and allogeneic bone marrow transplant recipients, respectively (Cesaro et al., 2018; Hirsch and Randhawa, 2019). Since specific antiviral agents and vaccines are not available, reducing immunosuppression is the current mainstay of therapy in order to regain control over BKPyV-replication (Cesaro et al., 2018; Hirsch and Randhawa, 2019).

However, this manoeuvre increases the risk of immunological injury such as allograft rejection and graft-versus-host disease. In our ongoing study to identify functionally and diagnostically relevant targets of BKPyV-specific antibody and T-cell responses (Binggeli et al., 2007; Cioni et al., 2016; Leboeuf et al., 2017), we noted that the small BKPyV-agnoprotein of 66 amino acids (aa) is abundantly expressed in the cytoplasm during the late viral life cycle *in vivo*, but largely ignored by the adaptive immunity (Leuenberger et al., 2007; Rinaldo et al., 1998). BKPyV-agnoprotein co-localizes with lipid droplets (LD) (Unterstab et al., 2010) and membranous structures of the ER (Unterstab et al., 2013). We now report that the BKPyV agnoprotein also targets mitochondria, disrupts the mitochondrial network and its membrane potential, and promotes mitophagy both in cell culture and in kidney allograft biopsies.

## Results

### BKPyV-Agnoprotein colocalizes with mitochondria and induces mitochondrial fragmentation

To investigate the function of BKPyV agnoprotein in the absence of LD, we noted that the N-terminal 20 aminoacid (aa) sequence shared some similarity with mitochondrial targeting sequences. To address the potential mitochondrial localisation, we infected primary human renal proximal tubular epithelial cells (RPTECs), a well-characterized model of BKPyV in renal allograft nephropathy (Bernhoff et al., 2008; Hirsch et al., 2016; Low et al., 2004). We compared infection of BKPyV-Dunlop (Dun-*AGN*) and the isogenic derivative Dun-*agn25D39E*. For this point mutant, binding to LD has been demonstrated to be abrogated after replacing the hydrophobic A25 and F39 with D and E, maintaining a central *α*-helix prediction, but abrogating its amphipathic character (Unterstab et al., 2010).

At 48 h post-infection (hpi) with wildtype Dun-*AGN* and mutant Dun-*agn25D39E* viruses, immunofluorescent staining detected both the early viral protein large T-antigen (LTag) and the late viral protein Vp1 capsid in the nucleus and agnoprotein in the cytoplasm (**Fig. S1**). Using the mitochondrial outer membrane protein Tom20 as a marker, specific colocalization of both the *AGN-*wildtype and *agn25D39E*-mutant protein was found, for the first time demonstrating agnoprotein colocalization to mitochondria (**Fig. 1A**). Of note, the mitochondria of Dun-*AGN-*infected RPTECs had lost the network-like pattern typical of uninfected cells and appeared in short, fragmented units (**Fig. 1A, supplementary Movie1**). In contrast, Dun-*agn25D39E*-infected RPTECs exhibited a regular mitochondrial network similar to uninfected cells (**Fig. 1B; supplementary Movie2**). Quantification of the mitochondrial morphology indicated a large excess of short mitochondrial fragments in Dun-*AGN-*infected cells, whereas mostly elongated mitochondria in a network-like pattern were seen in the Dun-*agn25D39E*-infected cells (**Fig. 1B**). Given these striking differences in mitochondrial phenotype, we examined whether or not the mutant *agn25D39E*-agnoprotein was still able to target the ER as reported for the wildtype agnoprotein (Unterstab et al., 2013). Confocal microscopy revealed that the *agn25D39E*-protein colocalized with the ER-marker calreticulin (**Fig. 1C**). However, whereas the mitochondrial colocalization of the *agn25D39E*-protein appeared in network strings, the ER-colocalization with calreticulin was patchy and reminiscent of the contact sites with the mitochondria-associated membranes (**Fig. 1C**). The patchy ER-colocalization pattern was independently confirmed using protein disulphide isomerase (PDI), another ER-marker protein (**Fig. S1C**). The results indicated that targeting of ER and mitochondria remained intact and implicated the amphipathic character of the central *α*-helix in disrupting the mitochondrial network.

**Fig. 1.**
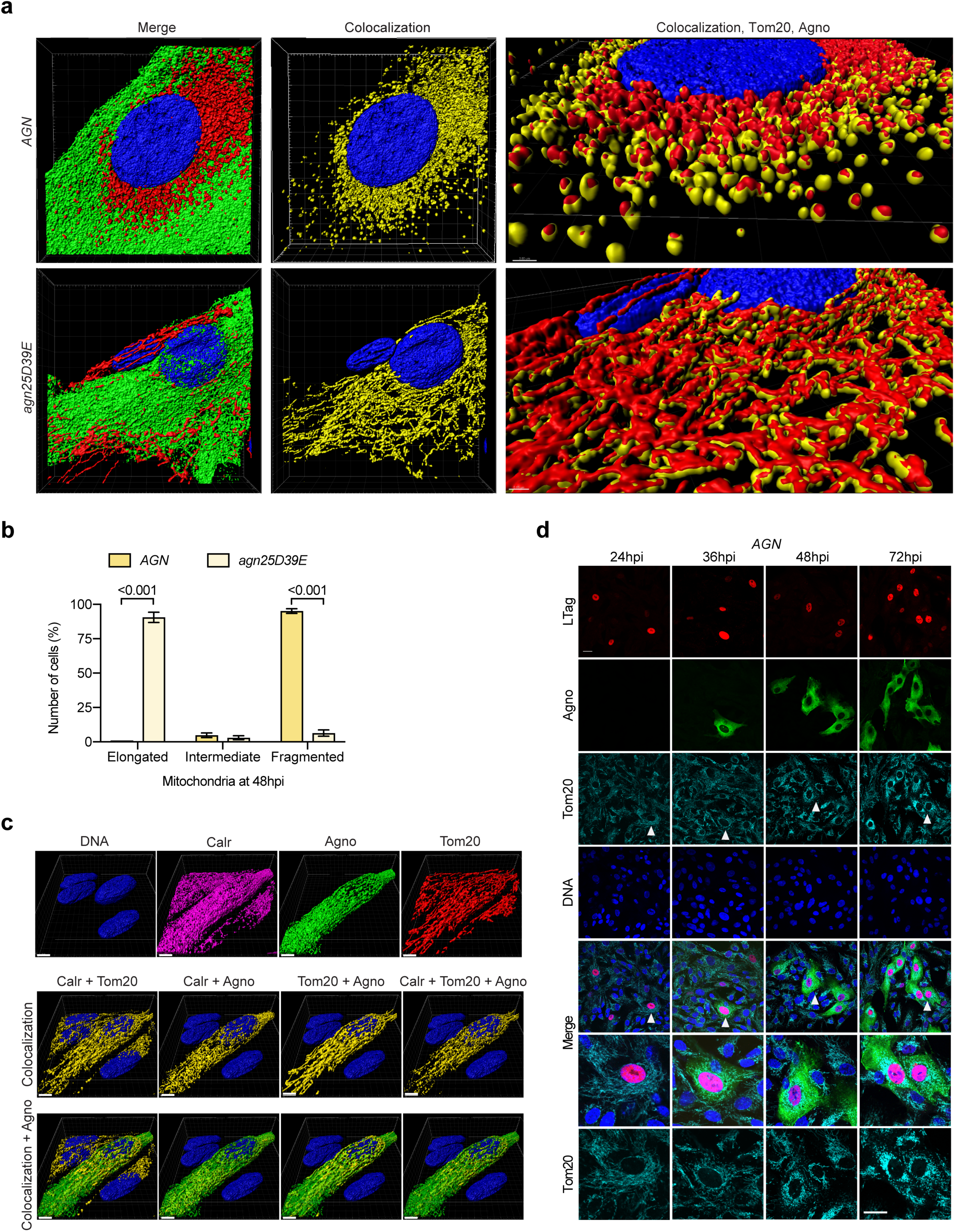
Agnoprotein colocalizes with mitochondria and induces mitochondrial fragmentation at late stage of BKPyV infection. Cells were infected with the indicated viral strains and fixed at 48 hpi as described in Materials & Methods. (a) Z-stacks of BKPyV Dun-*AGN* and Dun-*agn25D39E* infected cells, stained for Tom20 (red), agnoprotein (green), and DNA (blue). Colocalising voxels are shown in yellow. (b) Graph representing corresponding mitochondrial morphology. Quantification of six fields of two independent experiments using image J software (mean ± SD, 2-way ANOVA). (**Cc**) Z-stacks of BKPyV Dun-*agn25D39E* infected cells, stained for mitochondrial marker Tom20 (red), calreticulin as marker for the endoplasmic reticulum (magenta), agnoprotein (green), and DNA (blue). Colocalizing voxels are shown in yellow. Scale bar 5µm. (**d**) Confocal images of RPTECs infected with BKPyV-Dun-*AGN* at indicated time post infection. Cells were stained for LTag (red), agnoprotein (green), mitochondrial marker Tom20 (cyan), and DNA (blue). White arrows indicate cells magnified. Scale bar 20µm.

To correlate the strikingly altered mitochondrial morphology with the viral life cycle, we examined a time course of Dun-*AGN* infection demonstrating that expression of the early viral LTag at 24 h post-infection (hpi) had no effect on the mitochondrial network (**Fig. 1D**). After 36 hpi, agnoprotein expression started to appear in the cytoplasm, but mitochondrial fragmentation and perinuclear condensation became apparent only from 48 hpi onwards. At 72 hpi, Dun-*AGN*-infected cells could be readily identified by the fragmented mitochondrial network using Tom20 staining alone (**Fig. 1D**). Thus, wildtype and *agn25D39D*-agnoprotein were both able to localize to mitochondria and the ER, which in case of the wildtype led to mitochondrial fragmentation, while this was not observed for the Dun-*agn25D39D* mutant.

To further characterize the role of agnoprotein and its amphipathic helix, two additional isogenic BKPyV-derivatives were generated: Dun-*agn25L39L* encoding a mutant agnoprotein retaining the amphipathic character of the central *α*-helix, and Dun-*ATCagn*, in which the *ATG* start codon had been changed to *ATC* to create a “null”-agnoprotein virus. Ribbon models predicted that the overall secondary structure of *agn25D39E* and *agn25L39L* were similar to the *AGN*-encoded wildtype protein in having an N-terminal and a central helix, whereas the C-terminal structure was not predictable except for a short terminal *α*-helical tail (**Fig. 2A**). All BKPyV variants were found to proceed to the late viral replication phase, as evidenced by nuclear Vp1 staining (**Fig. 2B**). Similar to Dun-*AGN* wildtype infection, the Dun-*agn25L39L-*infected RPTECs exhibited a cytoplasmic agnoprotein distribution and mitochondrial fragmentation (**Fig. 2B**). In contrast, Dun-*ATCagn*-infected RPTECs retained intact mitochondrial networks similar to Dun-*agn25D39E,* but lacked agnoprotein expression as expected (**Fig. 2B**).

**Fig. 2.**
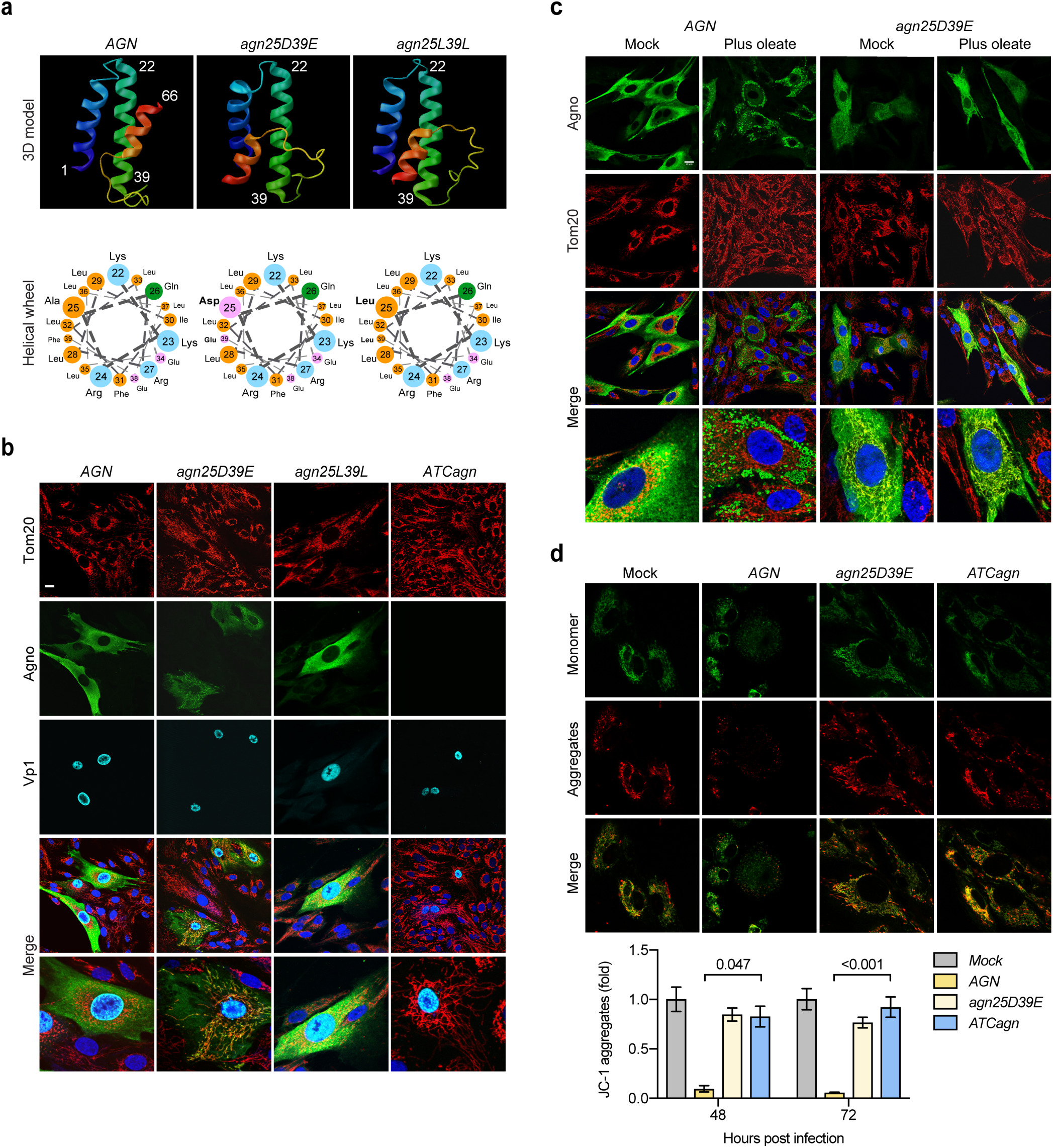
The amphipathic character of the central agnoprotein helix is required for mitochondrial fragmentation. (**a**) Three-dimensional ribbon model of each agnoprotein derivate as predicted with the Quark online algorithm (https://zhanglab.ccmb.med.umich.edu/I-TASSER/) and predicted amphipathic helical wheel of the agnoprotein aa22-39 (http://cti.itc.virginia.edu/~cmg/Demo/wheel/wheelApp.html). (b) Confocal images of RPTEC-infected with BKPyV Dun-*AGN* and its mutants, i.e., Dun-*agn25D39E*, *agn25L39L*, and Dun-*ATCagn*, respectively. Cells were fixed at 48 hpi and stained for mitochondrial marker Tom20 (red), agnoprotein (green), Vp1 (cyan), and DNA (blue). Scale bar 20µm. (c) Confocal images of RPTECs infected with BKPyV-Dun-*AGN* and BKPyV-Dun-*agn25D39E*, respectively, at 48 hpi. Cells were mock-treated or treated with 300µM Oleate at 24hpi. Cells were stained for Tom20 (red), agnoprotein (green), and DNA (blue). Scale bar 20µm. Live cell imaging using JC-1 dye (5µM) of -mock-infected or infected with BKPyV-Dun-*AGN* or its mutants, i.e., Dun-*agn25D39E*, and Dun-*ATCagn*, respectively at 48 hpi. Quantification of mitochondrial membrane potential (Ψ_m_) by measuring red fluorescent signal (JC-1 aggregates) with the Safire II plate reader of three independent experiments (mean ± SD, Kruskal-Wallis test).

Since LD-binding had been shown to require the amphipathic character of the central *α*-helix now implicated in mediating mitochondrial fragmentation (Unterstab et al., 2010), the effect of LD-formation on the mitochondrial network was investigated. To this end, RPTECs were infected with Dun-*AGN* or with Dun-*agn25D39E* and 300*μ*M oleate was added at 24 hpi.

Confocal microscopy of Dun-*AGN* at 48hpi revealed that agnoprotein was sequestered around LD, and that mitochondrial fragmentation of Dun-*AGN* infected cells was abrogated (**Fig. 2C**). In contrast, Dun-*agn25D39E-*infected RPTECs lacked LD-sequestering of the mutant agnoprotein despite the presence of cytoplasmic droplets presenting with punched-out holes (**Fig. 2C**) as reported (Unterstab et al., 2010). These results functionally linked the amphipathic helix with LD-binding and mitochondrial fragmentation in a competitive intracellular way. To investigate functional consequences of agnoprotein expression, we examined the mitochondrial membrane potential (MMP) using the (Ψ_m_)-dependent accumulation of JC-1 dye, whereby mitochondrial depolarization is indicated by a decrease in the red/green fluorescence intensity. At 48 hpi, Dun-*AGN*-infected cells exhibited fragmented mitochondria (green channel) and a significant decrease in MMP (red channel), whereas MMP changed little in Dun-*agn25D39E-* or Dun-*ATCagn-*infected RPTECs, or in mock-treated cells (**Fig. 2D**). Similarly, automated measurements of overall JC-1 red and green signals in cell cultures revealed significant MMP decreases of about 40% at 48 hpi in Dun-*AGN*-infected RPTECs compared to controls. The results indicated that infection with BKPyV expressing an agnoprotein with a central amphipathic helix was necessary for MMP-breakdown and network fragmentation in the late viral replication phase.

### Agnoprotein is sufficient for mitochondrial fragmentation and breakdown of the mitochondrial membrane potential and confers impaired innate immune signalling

To investigate whether or not agnoprotein expression alone is sufficient for mitochondrial fragmentation, expression vectors containing the full-length gene of wildtype agnoprotein or agnoprotein subdomains fused to monomeric enhanced green fluorescent protein (mEGFP) (Unterstab et al., 2010) were transfected into UTA6 cells (**Fig. 3A**). At 24 hours post-transfection (hpt) of the agno(1-66)mEGFP construct, GFP and agnoprotein overlapped in colocalization to fragmented mitochondria (**Fig. 3A, top row**). Similarly, mitochondrial colocalization and fragmentation was seen for agno(1-53)mEGFP lacking the C-terminal 13 aa of agnoprotein, but not for any of the other truncated agnoprotein constructs shown.

**Fig. 3.**
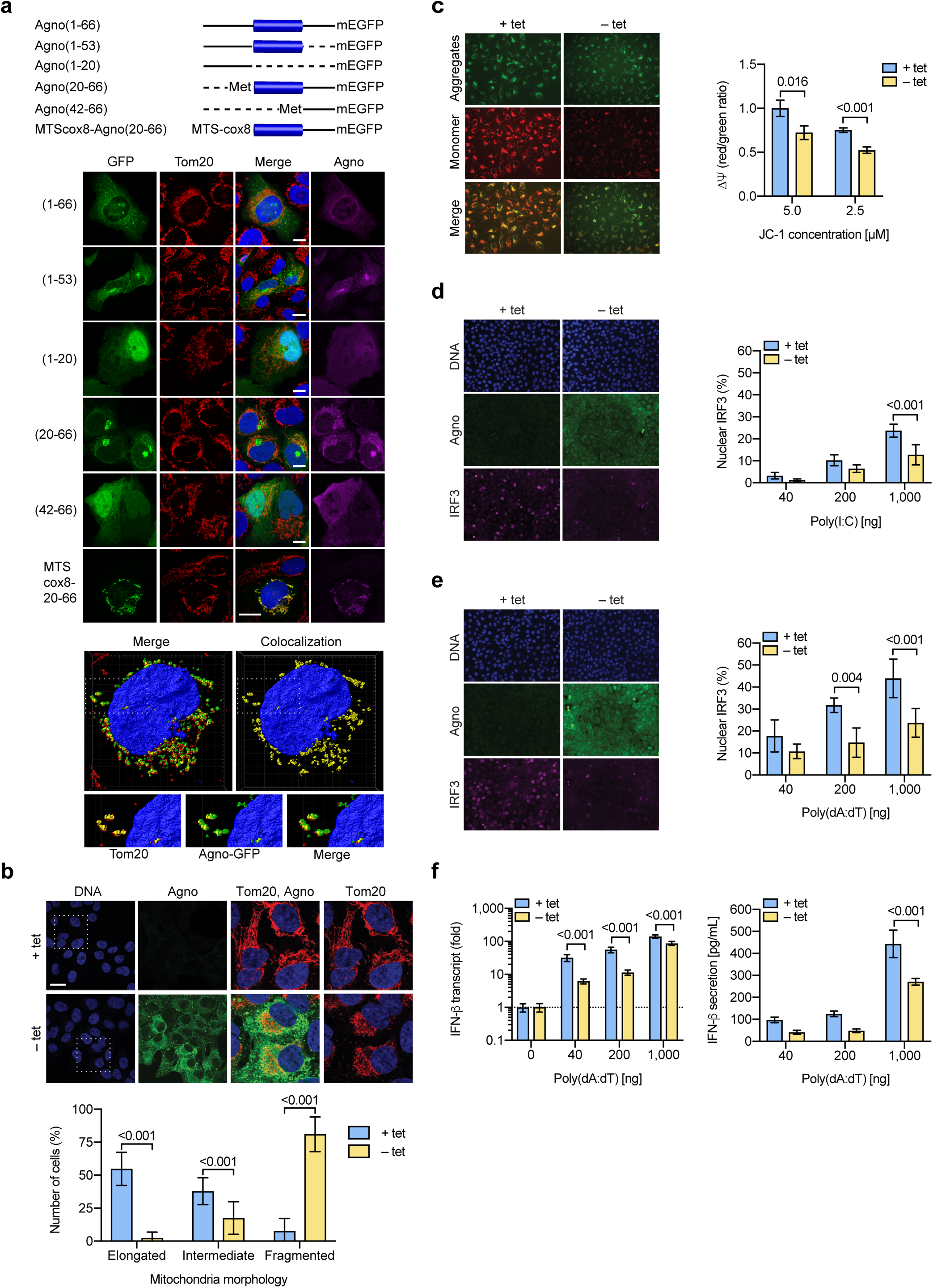
Agnoprotein is sufficient to mediate structural and functional alterations of mitochondrial network (a) Schematic presentation of agnoprotein-mEGFP fusion constructs transfected into UTA-6 cells. Retained aa indicated in parenthesis and presented as solid line, truncated parts presented as dotted line. The central amphipathic helix (aa 22–39) is shown as blue bar. Confocal images of transfected UTA-6 cells, transiently expressing the indicated agnoprotein-mEGFP fusion constructs were taken at 24 hpt. Immunofluorescent staining for Tom20 (red) and agnoprotein (magenta), GFP (green); DNA (blue). Scale bar 20 µm. Z-stacks were obtained from UTA-6 cells transfected with MTScox8-agno(20-66) at 24 hpt. Colocalizing voxels are shown in yellow, white dashed-lined rectangle indicating enlarged section. (b) UTA6-2C9 cells stably transfected with tetracycline (tet)-off inducible BKPyV agnoprotein were cultured for 24h in the presence or absence of tetracycline to suppress or induce BKPyV agnoprotein expression, respectively. Confocal images of cells stained for DNA (blue), agnoprotein (green), and Tom20 (red). White rectangle indicating enlarged section. Quantification of six fields using image-J of two independent experiments (mean ± SD; 2-way ANOVA) Ψ_m_ was evaluated using JC-1 dye and imaging of live cells using the signal ratio of aggregate (red)/ monomeric (green) normalized to UTA6-2C9 cells not expressing agnoprotein (tet+) versus cells expressing agnoprotein (tet-) using Mithras^2^ (mean ± SD,, unpaired parametric t-test). (c) Nuclear IRF3 translocation following poly(I:C) transfection was compared in UTA6-2C9 cells cultured for 24h in the presence or absence of tetracycline to suppress or induce BKPyV agnoprotein expression, respectively. Increasing amounts of rhodamine-labeled poly(I:C) was delivered to the cells via lipofection, cells were fixed at 4 hpt, stained for IRF3 (magenta), agnoprotein (green) and DNA (blue) (left images, 1000 ng poly(I:C), right panel quantification of nuclear IRF3 of six fields using Fiji (mean ± SD, Mann-Whitney). (d) Nuclear IRF3 translocation following poly(dA:dT) transfection was compared in UTA6-2C9 cells as described in D (left images, 1000 ng poly(dA:dT), right panel quantification of nuclear IRF3 (mean ± SD, Mann-Whitney). (e) Quantification of IFN-*β* mRNA and IFN-*β* secretion into cell culture supernatants following poly(dA:dT) stimulation of UTA6-2C9 cells, three experiments (mean ± SD, Mann-Whitney t-test).

Since the truncated agno(20-66)mEGFP had been previously demonstrated to be able to still colocalize with LD (Unterstab et al., 2010), we hypothesized that the N-terminal domain was necessary for mitochondrial targeting in order to mediate the breakdown of the mitochondrial network. Therefore, the N-terminal mitochondrial targeting sequence of cytochrome c oxidase (MTScox8) was fused to agno(20-66)-EGFP lacking its natural N-terminus. The results showed that agno(20-66)-EGFP was sufficient to induce mitochondrial fragmentation if mitochondrial targeting was provided by the MTScox8 sequences (**Fig. 3A, bottom row and extra panels; supplementary movie3**). For an independent confirmation, we examined UTA6-2C9 cells harbouring a wildtype full-length agnoprotein under the control of an inducible tetracycline (tet)-off promoter (Cioni et al., 2013). At 24 h in the absence of tet, agnoprotein was expressed in the cytoplasm and colocalized with fragmented mitochondria, whereas in the presence of tet concentrations suppressing agnoprotein expression, an intact mitochondrial network was seen (**Fig. 3B**). Importantly, inducing agnoprotein was associated with MMP disruption when examining aggregate (red) and monomeric (green) signals or when using automated JC-1 dye measurements of the cell cultures (**Fig. 3C**). Since mitochondria have a key role in innate immunity (Koshiba et al., 2011), UTA6-2C9 cells were cultured for 24h in the presence or absence of agnoprotein expression, and transfected with either poly(I:C)RNA or poly(dA:dT)DNA, both of which are potent inducers of the type-1-interferon expression via cytoplasmic phosphorylation and nuclear translocation of the interferon regulatory factor 3 (IRF3). At 4 hpt, the effect of agnoprotein expression on the nuclear localization of the IRF3 was examined. The results showed that in the presence of agnoprotein, nuclear IRF3 translocation was significantly reduced after transfecting poly(I:C) (**Fig. 3D**) or poly(dA:dT) (**Fig. 3E**). To functionally relate the agnoprotein-dependent differences in nuclear IRF-3 translocation, interferon (IFN)-*β* expression was quantified after transfection of increasing amounts of poly(dA:dT). The results indicated that agnoprotein expression resulted in a significant reduction of IFN-*β* transcripts and secreted protein levels (**Fig. 3F**). Together, the data indicated that the expression of the wildtype BKPyV-agnoprotein was necessary and sufficient to induce mitochondrial fragmentation, breakdown of MMP, and impairment of the innate immune sensing of transfected cytosolic DNA and RNA. BKPyV replication is inhibited by IFN-*β*.

To determine whether or not BKPyV replication is sensitive to IFN-*β*, RPTECs were pre-treated, which resulted in an IFN-*β* dose-dependent reduction of LTag-positive cells and supernatant BKPyV loads at 72 hpi (**Fig. 4A**). SDS/PAGE Immunoblot analysis demonstrated an increase of IFN-induced protein with tetratricopeptide repeats (IFIT) and a reduction of the viral capsid protein Vp1 and agnoprotein (**Fig. 4B**). A time course of IFN-*β* addition revealed maximal inhibitory effects before or at 2 hpi, but addition at 36 hpi still reduced the supernatant BKPyV loads by 50% (**Fig. 4B**). The reduction in supernatant BKPyV loads could be restored by type-I interferon blockade consisting of a cocktail of blocking antibodies against IFN-, IFN- and IFN-α alpha/beta receptor (**Fig. 4B**). Thus, BKPyV replication in primary human RPTECs was inhibited by IFN-*β*, but could be prevented by anti-IFN-blockade. To examine the role of agnoprotein in BKPyV replication, RPTECs were infected with equivalent infectious doses of Dun-*AGN* or the mutant derivatives Dun-*agn25D39E* and Dun-*ATCagn*. Although the mutant variants replicated, both had lower supernatant BKPyV loads at 72 hpi and a reduced number of LTag- and Vp1-positive cells compared to wildtype virus (**Fig. 4C**). To investigate if the reduced replication of Dun-*agn25D39E* involved innate immune activation, the TBK-1-inhibitor (Bx795) to prevent IRF3-phosphorylation, or the type-1 interferon-blocking cocktail was added at 24 hpi. The results demonstrated that TBK-1 inhibition or type-1 interferon blockade were able to partially rescue BKPyV Dun-*agn25D39E* replication (**Fig. 4C**). Since CoCl**_2_**-treatment has been described to induce functional hypoxia by disruption of the MMP (Jung and Kim, 2004), RPTECs were infected with Dun-*agn25D39E* and treated with 150 *μ*M or 300 *μ*M CoCl_2_ at 24 hpi. Partial mitochondrial fragmentation of BKPyV Dun-*agn25D39E* infected cells was seen together with increased supernatant viral loads at 72 hpi (**Fig. 4D**). Together, the results indicated that failure of the *agn25D39E* mutant protein to disrupt the MMP and mitochondrial network was associated with reduced replication, which involved innate immune activation and inhibition by type-1-interferon expression via mitochondrial signalling relays.

**Fig. 4.**
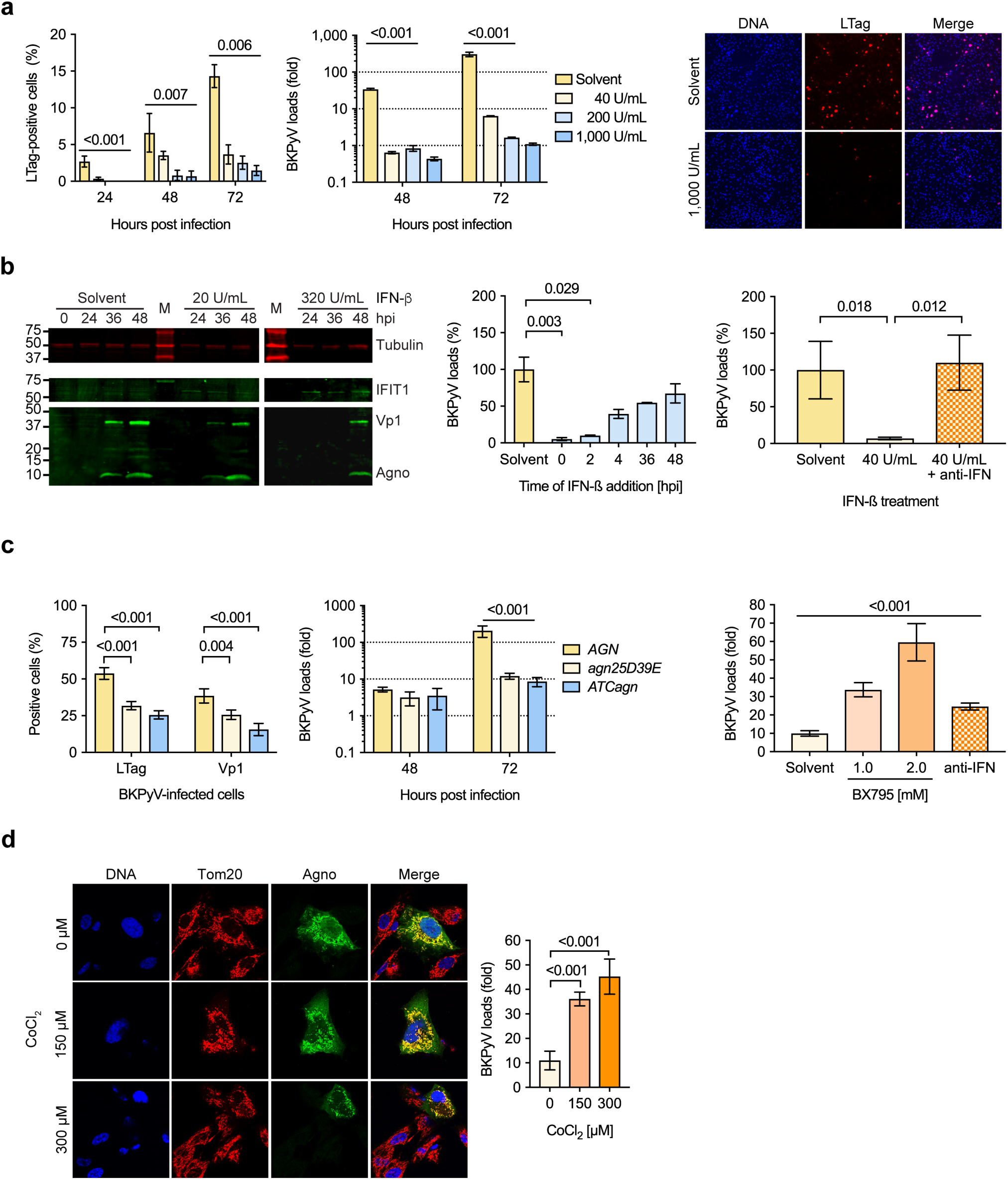
BKPyV replication in primary human RPTECs is sensitive to type-1 interferon. (a) RPTECs were treated with indicated concentrations of IFN-*β* or solvent overnight. LTag positive cells (left panel) and supernatant BKPyV loads (middle panel) were quantified at the indicated times, and a representative LTag staining of RPTECs at 72hpi is shown (right panel). (b) RPTECs were pre-treated with the indicated concentrations of IFN-*β* and expression of IFIT1 (*ISG56*) and BKPyV late viral proteins Vp1 and agnoprotein were analyzed at the indicated times post-infection by immunoblot analysis (left panel). RPTECs were treated before or at the indicated times post-infection with 200 U/mL IFN-*β*, and supernatant BKPyV loads were measured at 72 hpi (middle panel). BKPyV-infected RPTECs were treated with IFN-*β* in the presence or absence of anti-IFN consisting of antibodies blocking IFN-α, IFN-β and interferon alpha/beta receptor (right panel), and supernatant BKPyV loads were measured at 72 hpi (right panel; (right panel; duplicates, mean ± SD, unpaired parametric t-test). (c) RPTECs were infected with the indicated BKPyV variants (MOI = 1 by nuclear LTag staining of RPTCs) and the number of infected cells were quantified by immunofluorescence at 72 hpi (left panel) and supernatant BKPyV loads were measure at the indicated time post-infection (middle panel; triplicates, mean ± SD, 2-Way ANOVA). RPTECS were infected with BKPyV Dun-*agn25D39E*, the TBK-1 inhibitor BX795, antibodies blocking IFN-α β interferon alpha/beta receptor or solvent were added at 2 hpi, and supernatant BKPyV loads were measured at 72 hpi (right panel). (d) RPTECs were infected with BKPyV-Dun-*agn25D39E* and treated with the indicated concentrations of CoCl_2_ or solvent at 24hpi, and at 72 hpi, confocal microscop was performed for Tom20 (red), agnoprotein (green) and DNA (blue), and supernatant BKPyV loads quantified (right panel; triplicates, mean ± SD, unpaird parametric t-test).

Phosphorylation of the dynamin-related protein (Drp)1 at S616 by cell division kinase CDK1/cyclin B has been linked to its recruitment from the cytosol to mitochondria to physiologically mediate fission prior to mitosis and partitioning into daughter cells (**Fig. 5A, mock**). We therefore compared Drp1-S616 phosphorylation in BKPyV Dun-*AGN*, *agn25D39E*, and *ATCagn* infected RPTECs. Notably, we found an increased level of Drp1-S616 phosphorylation in BKPyV-infected Vp1-positive cells, in wildtype and *agn-*mutant viruses, hence independent of agnoprotein-mediated mitochondrial fragmentation (**Fig. 5A**). This indicated that Drp1-S616 phosphorylation was not sufficient for agnoprotein-mediated fragmentation of the mitochondrial network.

**Fig. 5.**
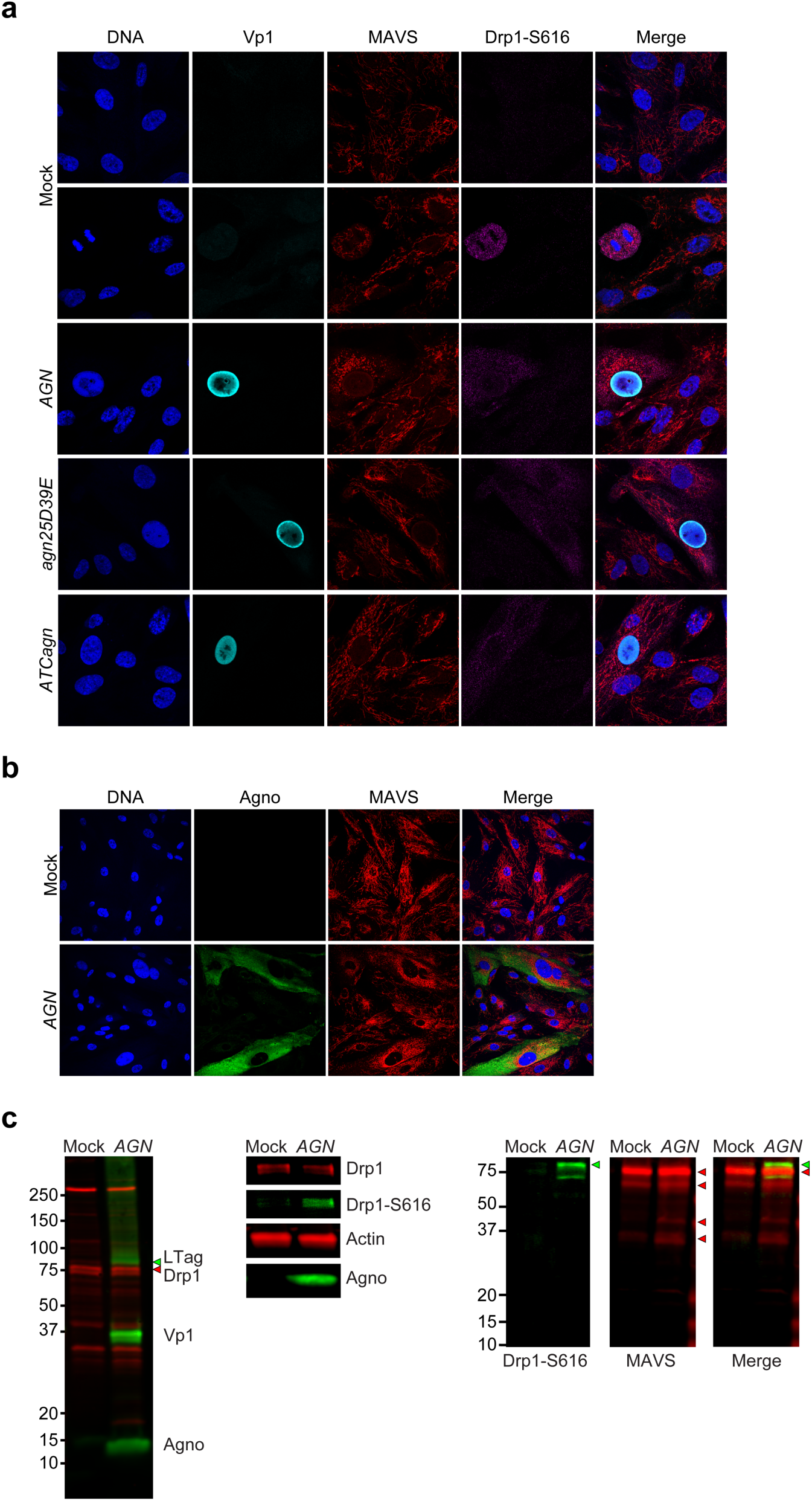
Drp-1 S616 phosphorylation and MAVS colocalization in BKPyV-infected RPTECs. (**a**) RPTECs were infected with BKPyV-Dun-*AGN*, Dun-*agn25D39E* or Dun-*ATCagn^null^* or mock-treated, and confocal microscopy was performed for Vp1 (cyan), Drp-1 S616 (magenta), MAVS (red), and DNA (blue). (b) RPTECs were infected with BKPyV-Dun-*AGN* or mock-treated, and at 72 hpi, confocal microscopy was performed for agnoprotein (green), MAVS (red), and DNA (blue). (c) RPTECs were infected with BKPyV-Dun-*AGN* or mock-treated and at 72hpi, cell lysates were prepared RIPA buffer and analysed by SDS/PAGE and immunoblotting using antibodies to DRP1-S616 (green triangle), MAVS (red triangle), and BKPyV agnoprotein (left panels) or DRP1-total protein and LTag, Vp1, and agnoprotein (right panel).

Since MAVS is an important sensor of cytosolic nucleic acids located on mitochondria, we examined its distribution in Dun-*AGN*-infected RPTECs by confocal microscopy. As shown, MAVS colocalized with both, intact as well as with agnoprotein-fragmented mitochondria in RPTECs (**Fig. 5B**). Similarly, MAVS also colocalized with fragmented mitochondria in UTA6-2C9 cells that expressed agnoprotein following tet-removal (data not shown). By SDS/PAGE and immunoblot analysis, no difference in Drp1 levels were seen, but a significant increase in Drp1-S616 phosphorylation was confirmed (**Fig. 5C, left panels**). To examine the possibility of MAVS degradation, cell extracts were analysed by SDS/PAGE/immunoblotting. The results revealed MAVS degradation with an increase in several smaller products in Dun-*AGN*-infected RPTECs compared to non-infected control cells (**Fig. 5C, right panels**).

Taken together, the results for the two mitochondria-associated proteins DRP-1 and MAVS indicated different effects of BKPyV-infection in RPTECs: no differences in DRP1-levels, but increased Drp1-S616 phosphorylation following BKPyV-infection independent of agnoprotein-mediated fragmentation, while MAVS remained colocalized to mitochondrial fragments and was degraded.

### Agnoprotein-disrupted mitochondria are targeted for p62/SQSTM1-mitophagy

Given the striking differences in mitochondrial structure by confocal microscopy, we examined BKPyV-Dun-*AGN* or Dun-*agn25D39E*-infected RPTECs at 72 hpi by transmission electron microscopy. The presence of viral particles in the nucleus served as a marker of the late viral life cycle equivalent to nuclear Vp1 staining. In the cytoplasm of Dun-*AGN*-infected RPTECs, small disrupted mitochondrial vesicles were seen, matching the results obtained by confocal microscopy, as well as several smaller dense multilaminar structures. In contrast, longer filamentous mitochondrial structures were seen in BKPyV Dun-*agn25D39E* (**Fig. 6A**) reflecting tangential cuts of the intact three-dimensional mitochondrial network.

**Fig. 6.**
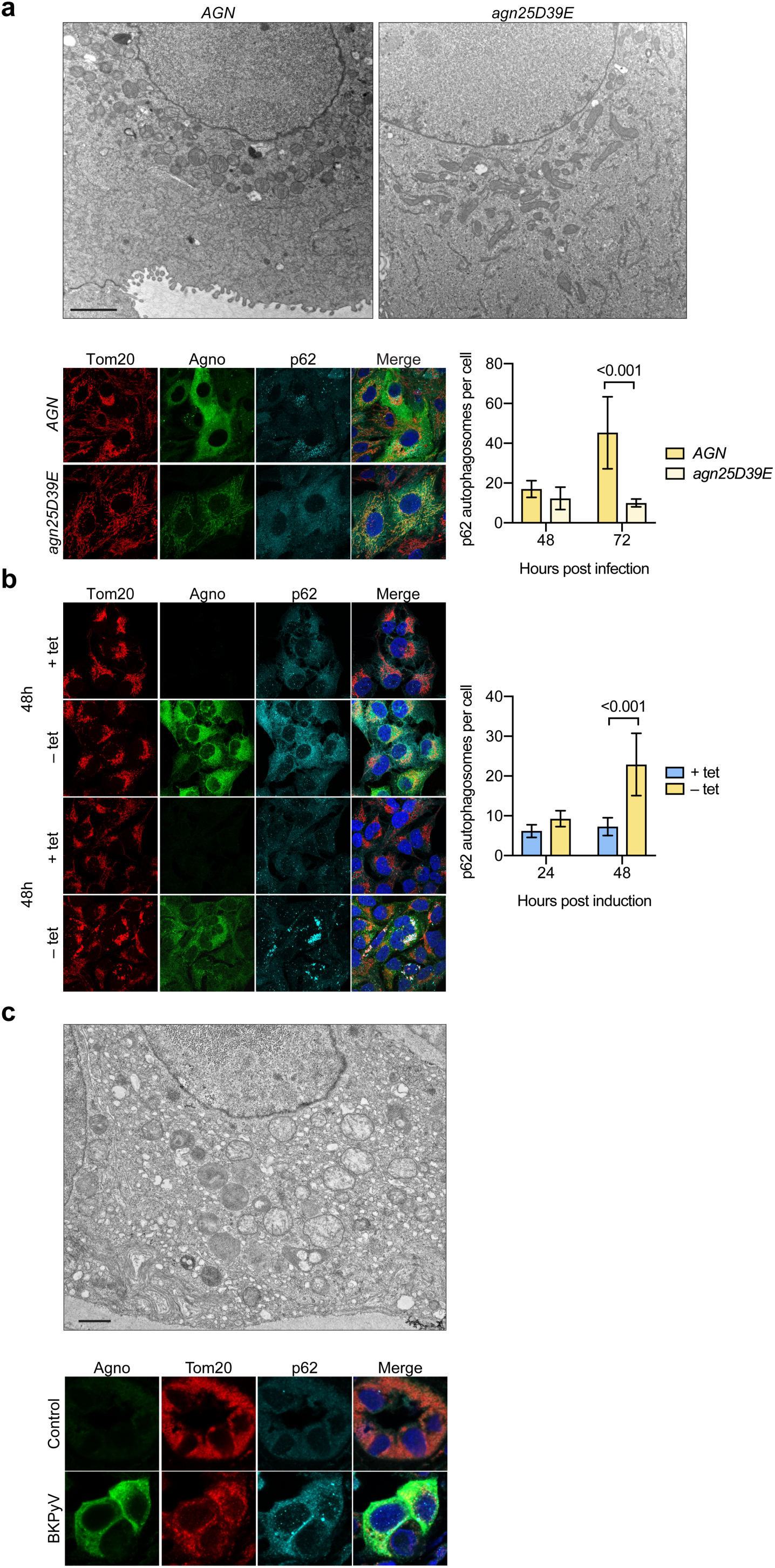
Agnoprotein-mediated mitochondrial fragmentation and p62/SQSTM1-autophagosomes is analysed by transmission electron microscopy and confocal microscopy of infected RPTECs and kidney transplant biopsy. (**a**) RPTECs were infected with BKPyV-Dun-*AGN* or BKPyV-Dun-*agn25D39E*. At 72hpi, cells were fixed and processed for TEM (top panels) or for confocal microscopy (bottom left panels) staining for Tom20 (red), agnoprotein (green), p62/SQSTM1 (cyan) and DNA (blue). p62/SQSTM1-positive autophagosomes of six fields were quantified using Fiji at 48 hpi and 72 hpi (bottom right panels; mean ± SD, 2-way ANOVA). (b) UTA6-2C9 cells were cultured for 24h or 48h in the presence or absence of tetracycline to suppress or induce BKPyV agnoprotein expression, respectively. At the indicated times, confocal microscopy (left panels) was performed after fixing and staining for Tom20 (red), agnoprotein (green), p62/SQSTM1 (cyan) and DNA (blue). p62/SQSTM1-positive autophagosomes were quantified as described in A at 24 h and 48 h (right panels; mean ± SD, 2-way ANOVA). (c) Tissue biopsies from kidney transplant patient suffering from BKPyV-associated nephropathy was studied by transmission electron microscopy (top panel) or confocal microscopy (bottom panel) staining for Tom20 (red), agnoprotein(green), p62/SQSTM1 (cyan), and DNA (blue).

Since damaged mitochondria with MMP breakdown may be targeted for autophagy, we investigated p62/SQSTM1 as a marker of mitochondrial autophagosomes. At 72 hpi, a significant increase in larger confluent punctate p62/SQSTM1-positive autophagosomes was observed in Dun-*AGN* infected RPTCs compared to the disperse cytoplasmic distribution in mutant Dun-*agn26D39E*-infected cells (**Fig. 6A**). Moreover, extensive cytoplasmic agglomeration and colocalization of p62/SQSTM1 and Tom20 was seen in Dun-*AGN-* infected RPTEC, which was rare in Dun-*agn25D39E*-infected cells (**Fig. S2**). Similarly, agnoprotein-dependent p62/SQSTM1-accumulation was seen in UTA6-2C9 cells at 24h post-induction, which increased until 48h indicating that agnoprotein-induced breakdown of the mitochondrial membrane potential and network fragmentation was followed by p62/SQSTM1-positive autophagosome formation (**Fig. 6B**).

To investigate whether or not similar changes could be observed in BKPyV-associated nephropathy in kidney transplant patients, allograft biopsy samples were analysed. Indeed, transmission electron microscopy revealed small disrupted mitochondria in the cytoplasm of tubular epithelial cells having viral particles in the nuclei (**Fig. 6C**). Immunofluorescent staining of kidney allograft biopsies and confocal microscopy for agnoprotein, Tom20 and p62/SQSTM1 revealed fragmented mitochondria and autophagosomes (**Fig. 6C**). Together, the data indicated BKPyV replication in RTECS in cell culture or in renal allograft nephropathy was associated with mitochondrial fragmentation and p62/SQSTM1-accumulation implicated in mitophagy (Johansen and Lamark, 2011).

To investigate autophagic flux as a dynamic marker of p62/SQSTM1-mitophagy, the expression levels of LC3-I and its activated lapidated derivative LC3-II were examined in UTA6-2C9 cells by immunoblotting. In untreated cells, virtually only LC3-I was detected, but in the presence of the lysosomal protease inhibitors pepstatin-A1/E64d, LC3-I increased and LC3-II became apparent as expected for inhibition of the steady-state autophagic flux (**Fig. 7A**). Following agnoprotein expression, LC3-I and the derivative LC3-II levels were also increased, and increased further in the presence of pepstatin-A1/E64d indicating that agnoprotein expression increased the autophagic flux, which could be partly blocked by protease inhibitors (**Fig. 7A**). Similar to agnoprotein expression alone, treatment with carbonyl cyanide m-chlorophenylhydrazone (CCCP) known to chemically disrupt the MMP and to induce PINK-Parkin-dependent mitophagy (Narendra et al., 2008) resulted in an increase in LC3-I and -II, and which further increased in the presence of pepstatin-A1/E64d (**Fig. 7A**). However, agnoprotein expression followed by CCCP treatment did not result in a further increase of LC3-I/II levels, but rather increased the relative LC3-II proportion at an overall lower level suggesting that the autophagic flux induced by agnoprotein was further maximized by CCCP treatment. To investigate the role of parkin in this process, UTA6-2C9 cells were transfected with a yellow fluorescent protein (YFP)-Parkin expression construct and analysed by confocal microscopy. In YFP-transfected UTA6-2C9 cells not expressing agnoprotein (tet+), little co-localization with fragmented mitochondria was observed.

**Fig. 7.**
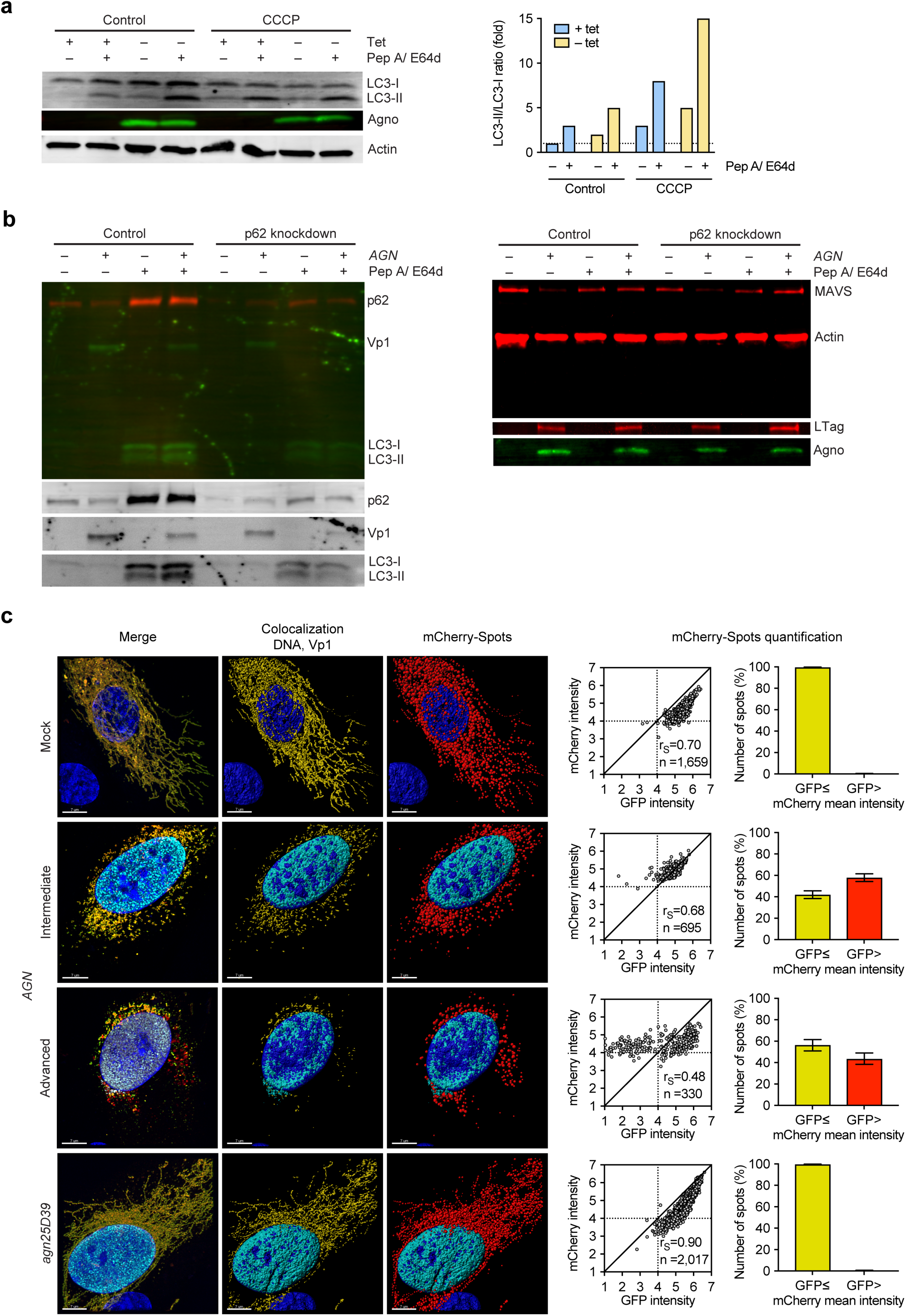
Agnoprotein mediates p62/SQSTM1-dependent autophagic flux and mitophagy. (a) UTA6-2C9 cells were cultured for 48 h in the presence or absence of tetracycline to suppress or induce BKPyV agnoprotein expression, respectively. At 24 h post-induction, pepstatin-A1/E64d (10µg/mL) and CCCP (10µM) was added as indicated and cell extracts were prepared and analysed by immunoblotting as described in Materials & Methods, using RIPA buffer, for LC3-I and II, agnoprotein expression and actin. LC3-II/I ratio were normalized using untreated cells without agnoprotein (+tet) expression as reference. (b) RPTECs transfected with siRNA-p62 (p62 knockdown) or control were infected with BKPyV Dun-*AGN* (indicated as *AGN*). Pepstatin-A1/E64d (10µg/mL) was added at 48 hpi, when indicated, 2.0×10^4^ cells were harvested and lysed per 10µL Laemmli Sample Buffer and analysed by immunoblotting as described in Material & Methods for p62/SQSTM1, BKPyV Vp1, and LC3-I and -II expression (on 0.22µm PVDF membrane, left panel) or MAVS, actin, BKPyV LTag, and agnoprotein (on 0.45 PVDF-FL membrane, right panel). (c) RPTECs transfected with mCherry-GFP-OMP25TM tandem tag mitophagy reporter were infected with BKPyV Dun-*AGN* or BKPyV-Dun-*agn25D39E*. At 72 hpi, cells were fixed and stained for Vp1 and DNA. Colocalization of mCherry and GFP is shown (yellow), Vp1 (cyan), and DNA (blue). Z-stacks were acquired and analyzed in IMARIS, the mCherry signal was transformed into countable spots (according voxel intensity). The GFP and mCherry mean intensity within the spots were quantified (bars represent mean ± 95% CI, Wilson/Brown method).

Following the addition of CCCP, strong cytoplasmic YFP-parkin positive agglomerates developed in the cytoplasm (**Fig. S3**). In UTA6-2C9 cells expressing agnoprotein (tet-), the YFP-Parkin signals appeared displaced by agnoprotein-Tom20 colocalizing structures accumulating in the perinuclear cytoplasm. Together, the data suggested that CCCP- and agnoprotein-induced mitophagy differed in being Parkin-dependent and -independent, respectively.

To independently investigate the role of p62/SQSTM1 in autophagic flux following BKPyV infection of RPTECs, LC3 was analysed in RPTEC with or without siRNA-p62 knockdown. Immunoblotting of the siRNA-p62 knockdown RPTECs revealed a reduction of p62/SQSTM1 levels to approximately 20 percent of the control RPTECs (**Fig. 7B, left panel**). Upon pepstatin-A1/E64d treatment, p62/SQSTM1 levels as well as the levels of LC3-I and -II increased in the control cells, but not to the same extent in the siRNA-p62 knockdown cells, indicating that the decrease in p62/SQSTM1 levels was associated with lower LC3-II formation and reduced steady-state autophagic flux (**Fig. 7B, left panel**). Following Dun-*AGN*-infection and pepstatin-A1/E64 treatment, LC3-I and -II levels increased, but the overall levels were lower in the siRNA-p62 knock-down cells, and the LC3-II/-I ratio was decreased as well as the Vp1 levels (**Fig. 7B, left panel**). To investigate the impact of the siRNA-p62 knockdown, whole cell extracts were prepared and analysed by SDS/PAGE-immunoblotting for MAVS levels in infected and uninfected RPTECs. The results revealed that MAVS levels were not affected by siRNA-p62-knockdown, but decreased upon BKPyV-AGN-infection, which could be partly inhibited in the presence of the lysosomal protease inhibitors pepstatin-A1/E64d. Together, the data suggested that BKPyV Dun-*AGN* infection of RPTECs was associated with an increased p62/SQSTM1-dependent autophagic flux, which involved MAVS degradation, and could be reduced by siRNA-p62 knock-down and pepstatin-A1/E64d treatment.

To further investigate mitophagy in BKPyV infection, the tandem tag mitophagy reporter mCherry-mEGFP-OMP25TM carrying the transmembrane domain (TM of OMP25) for targeting to the mitochondrial outer membrane (Bhujabal et al., 2017) was transfected into RPTECs infected with Dun-*AGN* or Dun-*agn25D39E*. Quantifying the red and green signals in z-stacks of Vp1-expressing cells following confocal microscopy revealed that the number of mitochondrial signals with mCherry red signals exceeding GFP green signal were higher in Dun-*AGN*-infected RPTECs and increased as the number of residual mitochondrial fragments progressively decreased compared to Dun-*agn25D39E*-infected cells or mock-treated controls (**Fig. 7C**). The data indicated that the mitochondrial tandem tag reporter was associated with the mitochondrial network in RPTECs infected with Dun-*AGN* or Dun-*agn25D39E*, but was progressively disrupted and targeted to the acidic environment of autophagosomes in the former. Together, the data supported the notion that BKPyV Dun-*AGN* infection of RPTECs was associated with increased mitophagy, which was not observed for infection with the mutant Dun-*agn25D39E*.

### Mitochondrial colocalization and fragmentation of agnoproteins is conserved among BKPyV, JCPyV, and SV40

To investigate the impact of agnoprotein expression in another BKPyV strain, infection of RPTECs was studied using a well-characterized, yet slowly replicating BKPyV-WW(1.4) strain carrying an archetype NCCR (Bethge et al., 2015; Gosert et al., 2008). Similar to Dun-*AGN*, the BKPyV-WW(1.4) strain showed agnoprotein expression associated with mitochondrial fragmentation (**Fig. 8A**). Since agnoprotein homologues have been identified in the human polyomavirus JCPyV, we examined SVG-A cells infected with JCPyV-Mad4 strain (Gosert et al., 2010). Colocalization of JCPyV-agnoprotein with Tom20 and mitochondrial fragmentation was observed (**Fig. 8B**; **Fig. S5**). Similarly, fragmented mitochondria were seen in CV-1 cells infected with the monkey virus SV40 showing Vp2/3-expression as a marker of late viral replication phase (**Fig. 8C**) or, since SV40-agnoprotein antibodies were not available, in cells showing cytoplasmic staining positive for the partly cross-reacting JCPyV-antibody (**Fig. 8C**). Together, the data indicated that the function of agnoprotein to support viral replication by mitochondrial fragmentation is conserved among renotropic polyomaviruses.

**Fig. 8.**
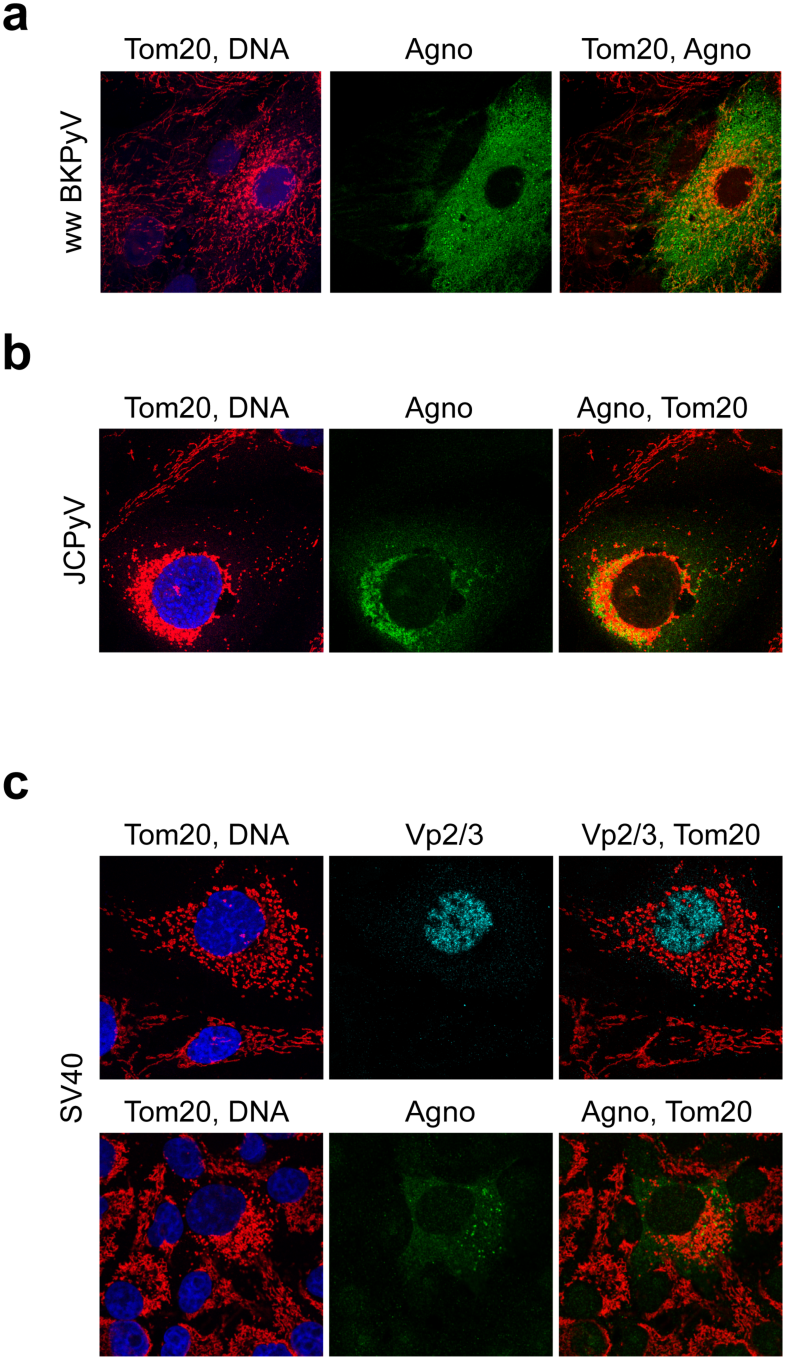
Mitochondrial fragmentation is a conserved feature observed in archetype BKPyV strains, JCPyV and SV40. (**a**) RPTECs were infected with BKPyV-WW(1.4) carrying an archetype non-coding control region and confocal microscopy was performed at 6 dpi after staining for Tom20 (red), agnoprotein (green), and DNA (blue). (b) SVG-A cells were infected with JCPyV-*Mad4* carrying an archetype non-coding control region and confocal microscopy was performed at 72 hpi after staining for Tom20 (red), anti-JCPyV agnoprotein (green), and DNA (blue). (c) CV-1 cells were infected with SV40 X and confocal microscopy was performed at 48 hpi after staining for Tom20 (red), Vp2/3 (blue) or cross-reacting antisera raised against JCPyV agnoprotein (green).

## Discussion

In the last decade, significant knowledge has been accumulated on how cytoplasmic sensing of viral infections is achieved by the innate immune system (Takeuchi and Akira, 2010; Zevini et al., 2017) and how this crucial first line of defense is subverted to allow for transient or persistent immune escape of acute and chronic viral infections, respectively (Garcia-Sastre, 2017; Goubau et al., 2013). Despite a high infection rate (Virgin et al., 2009) and evidence of immune evasion in the general population (Egli et al., 2009), comparatively little is known about relevant mechanisms operating in small non-enveloped DNA viruses such as human polyomaviruses. In this study, we report that the small BKPyV-encoded agnoprotein of 66aa facilitates polyomavirus replication by disrupting the mitochondrial network and its membrane potential during the late phase of the viral replication cycle.

Thereby, BKPyV replication is able to evade cytosolic innate immune sensing and to promote p62/SQSTM1-assisted mitophagy in this critical phase of viral progeny accumulation, during which DNA damage cumulates (Hein et al., 2009) and abundant viral DNA genomes and RNA transcripts exhaust in the infected host cell (Funk, 2008; Funk et al., 2006). Due to the precise timing to the critical late viral replication phase (Bernhoff et al., 2008; Low et al., 2004), this window of immune evasion is equally opened during *de novo* infection or reactivation permitting cell-to-cell spread below the radar of the immune system.

Transfection experiments using the entire agnoprotein or different subdomains fused to the reporter mEGFP as well as the tet-off inducible agnoprotein expression indicate that agnoprotein is necessary and sufficient, using its amino-terminal and central helix for mitochondrial targeting and mitochondrial disruption, respectively. As expected from the key role of mitochondria in relaying innate immune sensing, the agnoprotein-mediated disruption of the MMP was associated with significantly reduced IRF3 translocation into the cell nucleus as well as IFN-*β* transcript and protein expression following poly(I:C)RNA or poly(dA:dT)DNA stimulation. Importantly, BKPyV viral variants either lacking agnoprotein expression due to a start codon mutation (Dun-*ATCagn^null^*) or carrying mutations abrogating the amphipathic character of the central helix (Dun-*agn25D39E*) showed intact mitochondrial networks and little mitophagy, but significantly impaired replication compared to the wildtype strain Dun-*AGN*. The lower replication rate of the mutant Dun-*agn25D39E* could be partially reversed on three levels of the innate immune response, namely by CoCl_2_ treatment disrupting the mitochondrial membrane potential (Jung and Kim, 2004), by BX795 inhibiting the downstream signalling kinase TBK-1, or by type-1 interferon blockade. Although BKPyV as a DNA-virus would be expected to preferentially be sensed via cGAS/STING, increasing data indicate a close interaction between DNA- and RNA-sensing pathway coupling MAVS- and STING-activation on mitochondria and endoplasmic reticulum to TBK-1 activation downstream including through RNA-polymerase III transcription of cytoplasmic DNA-fragments (Chiu et al., 2009; Zevini et al., 2017). Although we can presently not exclude a more direct role of agnoprotein on STING signalling through MMP-breakdown or through its ER-co-localization, our results demonstrate that agnoprotein inhibits innate immune responses to both poly(I:C)-RNA and poly(dA:dT)-DNA. Notably, experimental MMP breakdown by CCCP has been shown to suppress STING signalling, further merging the cytosolic RNA and DNA-triggered responses (Kwon et al., 2017). Thus, we conclude that BKPyV-agnoprotein mediates MMP breakdown and antiviral immune subversion similar to other DNA and RNA viruses (Koshiba et al., 2011).

Our study also reveals for the first time a critical of mitophagy in polyomavirus biology. Mitophagy is known to assist in the disposal of irreversibly damaged mitochondrial fragments. The pepstatin-A1/E64d protease inhibition and siRNA-knockdown experiments indicate that agnoprotein increases the p62/SQSTM1-dependent steady-state autophagic flux via LC3-II lipidation in a fashion similar to the one observed following the CCCP-induced mitochondrial membrane potential breakdown (Johansen and Lamark, 2011). However, whereas CCCP-induced mitophagy involves PINK-Parkin pathway (Bhujabal et al., 2017), our results suggest that agnoprotein rather induced parkin-independent mitophagy. Further evidence for increased autophagic flux in BKPyV infected cells was obtained using the tandem tag mitophagy reporter mCherry-mEGFP-OMP25TM revealing acidification in mitophagosomes of BKPyV-*AGN*-infected RPTECs, which is not observed in *agn25E39D*-mutant infected RPTECs. Notably, pepstatin-A1/E64d protease inhibition and p62-siRNA knockdown, both of which act downstream of the agnoprotein-mediated impaired innate immune sensing and reduced interferon production, were associated with the reduced Vp1-protein levels in BKPyV Dun-*AGN*-infected RPTECs. This novel observation suggests that the autophagic flux may be relevant for the effective biosynthesis in the late viral replication phase following the functional and structural loss of the mitochondrial power plants (Forbes, 2016).

Innate immune sensors have been characterized in primary human RPTECs and in kidney biopsies by transcriptional profiling identifying antiviral, proinflammatory and proapoptotic responses (Heutinck et al., 2012; Ribeiro et al., 2012). Other studies reported that BKPyV infection of RPTECs can occur without induction of innate immune sensors by as yet unknown mechanisms (Abend et al., 2010; An et al., 2019; de Kort et al., 2017). However, these studies failed to reveal inhibition of BKPyV replication by type-1 interferons, which is in contrast to our results for BKPyV shown here and those recently reported for JCPyV (Assetta et al., 2016).

Our results also shed new light on previous reports on BKPyV agnoprotein and the closely related JCPyV and SV40 homologues (Gerits and Moens, 2012; Saribas et al., 2018): These include facilitating polyomavirus replication (Ng et al., 1985), increasing viral late phase production (Carswell et al., 1986) and plaques size (Hou-Jong et al., 1987), acting as egress factor (Panou et al., 2018) or as viroporin (Suzuki et al., 2010). We demonstrate that mitochondrial fragmentation occurs not only for BKPyV strains carrying an archetype NCCR, but also for the human JCPyV or the monkey SV40 supporting the view that this dramatic agnoprotein-mediated function is evolutionary conserved across different species, cell types and hosts.

Importantly, our results are confirmed in biopsies obtained from kidney transplant patients providing independent evidence of mitochondrial fragmentation and p62/SQSTM1-mitophagy *in vivo*. These data suggest that the combined effects of immune escape, mitophagy, and viral spread are operating not only in a relevant primary human cell culture model of RPTECs, but also in one of the currently most challenging pathologies affecting kidney transplantation (Ramos et al., 2009). Intriguingly, the BKPyV agnoprotein-induced immune subversion and p62/SQSTM1-mitophagy may contribute to the earlier reported paradox of abundant cytoplasmic expression *in vivo* and the low agnoprotein-specific antibody and T-cell response (Leuenberger et al., 2007), and provide a new twist to BKPyV-promoting hypoxic mechanisms following renal ischemia/reperfusion injury (Atencio et al., 1993; Fishman, 2002; Hirsch et al., 2006).

Immune evasion inactivating the cytosolic sensing platforms has been described for other human viruses (Khan et al., 2015) indicating that a huge variety of different steps up- and down-stream of mitochondria and associated ER-membranes can be perturbed in a virus-specific manner (Ding et al., 2017). However, the small BKPyV agnoprotein is unique by compactly and effectively combining several functional and structural properties ascribed to different other viral regulators. Thus, the matrix protein (M-protein) of parainfluenza-3 has been reported to translocate to mitochondria, where it induces mitophagy via LC3-II in a PINK/Parkin-independent manner (Ding et al., 2017). However, the M-protein is a structural protein, and according to the current model acts early following entry into host cells and requires piggy-back import via the mitochondrial elongation factor TUFM (Ding et al., 2017), while our study indicates that the non-structural agnoprotein targets mitochondria directly. This property of BKPyV agnoprotein seems to be shared with the influenza PB-F1 protein, which has been reported to translocate via the TOM40 channel (Gibbs et al., 2003). Unlike BKPyV, HCV and some flaviviruses replicating in the cytoplasm appear to induce mitochondrial fragmentation and mitophagy in a PINK-Parkin-dependent fashion (Gou et al., 2017). Although structurally unrelated to agnoprotein, the 247aa-long cytomegalovirus (CMV) US9 glycoprotein appears to suppress both MAVS and STING pathways, TBK-1 activation and IFN-*β* expression by disrupting the mitochondrial membrane potential in the late CMV replication phase (Choi et al., 2018; Mandic et al., 2009). Given the association of agnoprotein with LD, we were intrigued by the similarity to the antiviral host cell protein viperin, which is expressed following RNA- and DNA-virus infections (Gizzi et al., 2018; Hee and Cresswell, 2017). The antiviral effects of viperin have been attributed to a perplexing variety of functions including interference with lipid metabolism and formation of detergent-resistant lipid rafts at the sites of influenza budding (Wang et al., 2007). Similar to agnoprotein, Viperin contains a N-terminal amphipathic helix required for adsorbing to the cytosolic face of LD- and ER-membranes (Hinson and Cresswell, 2009a, b). Notably, viperin may facilitate viral replication (Hee and Cresswell, 2017), when re-directed to mitochondria by the CMV-encoded Bax-specific inhibitor viral mitochondria-localized inhibitor of apoptosis (vMIA) carrying a mitochondrial targeting domain (Cam et al., 2010; Seo et al., 2011). Thus, the CMV-hijacked vMIA-viperin complex shares properties accommodated in only 66aa of BKPyV-agnoprotein.

Although our study indicates that agnoprotein is both necessary and sufficient, it is presently unclear, whether or not the agnoprotein-mediated MMP breakdown and mitochondrial fragmentation require the direct interaction with cellular proteins other than the mitochondrial import machinery. Given the abundance of BKPyV agnoprotein in the cytoplasm and the clearly differing colocalization patterns to ER and to mitochondria, further studies need to be carefully conducted in order to avoid misleading artefacts. For the Bcl-2 family of proteins, a hierarchy of interactome complexes has emerged over the last decade (Bleicken et al., 2017; Edlich et al., 2011). Moreover, for the related JCPyV-agnoprotein, a strong tendency to form multimeric aggregates *in vitro* has been reported (Sami Saribas et al., 2013), which undoubtedly reduces the specificity of otherwise straight-forward pull-down approaches.

Presently, we favour a minimal model in which mitochondrial targeting of N-terminal domain of agnoprotein allows for embedding the central amphipathic helix into the outer leaflet of the outer mitochondrial membrane similar to other amphipathic proteins, where it remains available for LD-binding (**Fig.9**), and that this simple plug-and-play embedment suffices to progressively leak protons causing loss of membrane potential, preventing TBK-1 activation, and targeting for p62/SQSTM1 mitophagy (Jarsch et al., 2016; McMahon and Gallop, 2005; Shen et al., 2012).

**Fig. 9.**
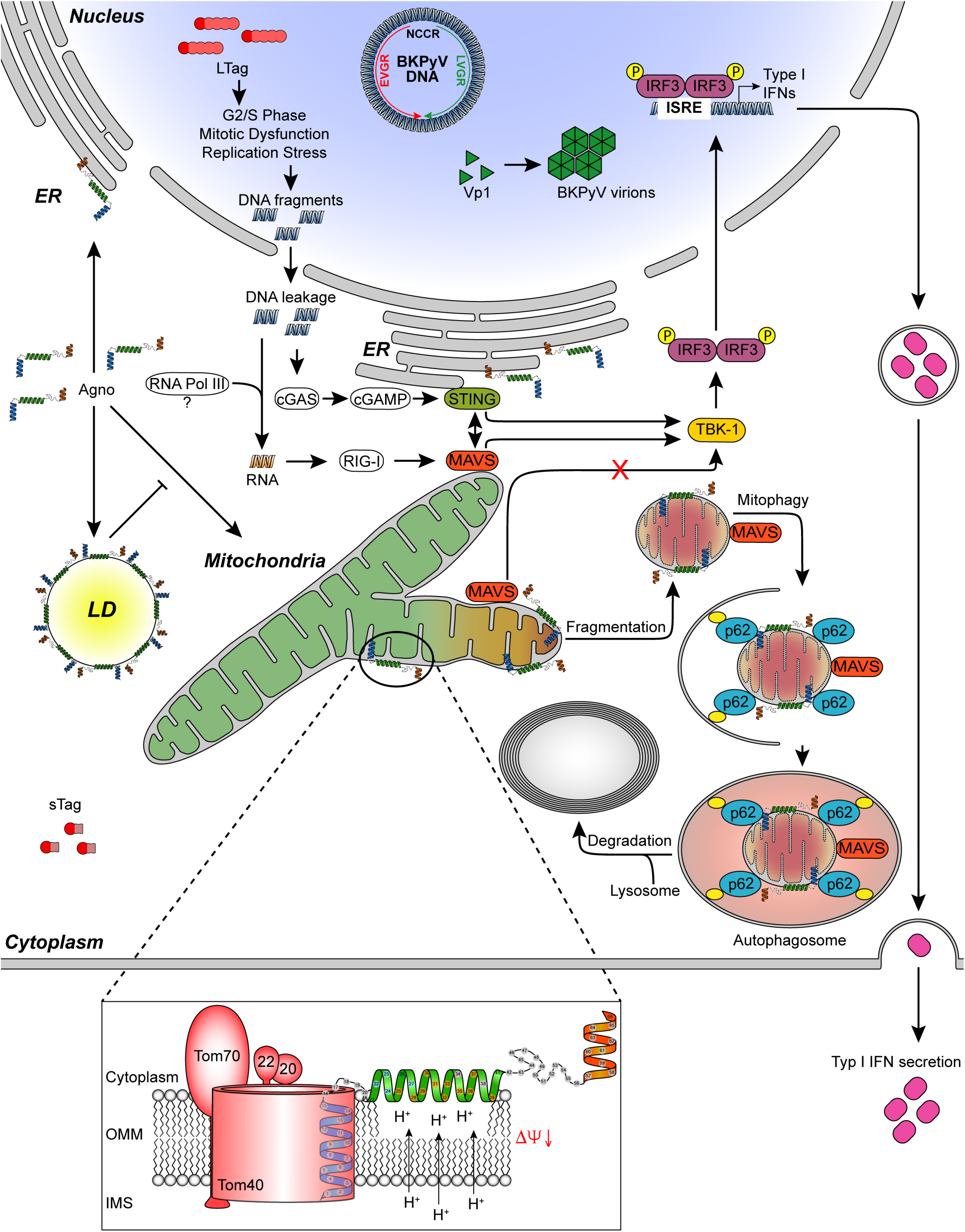
Working model of BKPyV agnoprotein mediating innate immune evasion by mitochondrial membrane breakdown, network fragmentation and mitophagy.

Thus, the combined data of our study provide important novel insights in how BKPyV-replication evades innate immune sensing in the critical late phase of the viral life cycle through the combined effects of the small accessory agnoprotein by at least three complementary effects: i) inactivating immune sensing and the inhibitory effects of interferon-*β* expression by breakdown of the mitochondrial membrane potential; ii) enhancing the supply of biosynthetic building blocks via increased autophagic flux, and iii) facilitating viral release of nuclear virions by fragmenting the nucleus-surrounding mitochondrial network. Finally, our observations suggest that BKPyV agnoprotein is not only important for immune escape and urinary shedding in healthy immunocompetent hosts, but may also facilitate the rapid cell-to-cell spread in the renal tubulus progressing to BKPyV-nephropathy in kidney transplant patients. These insights may help to design novel antiviral and immunization strategies and permit identifying novel markers readily distinguishing BKPyV nephropathy from allograft rejection (Hirsch and Randhawa, 2019).

## Materials and Methods

### Reagents

To induce LDs, culture medium was supplemented with 300 μ Aldrich) bound to essentially fatty acid free bovine albumin (A6003; Sigma-Aldrich) as described (Unterstab et al., 2010). Poly(I:C) (HMW) Rhodamine (tlrl-picr, InvivoGen) was transfected with ViaFect™ Transfection Reagent (E4982, Promega) at a reagent: DNA ratio of 3:1 according to manufacturer’s instructions or with Poly(I:C) (LMW)/LyoVec™ (tlrl-picwlv, InvivoGen). The treatment was administered as indicated. Poly(dA:dT) (tlrl-patn, InvivoGen), Rhodamine (tlrl-patrh, InvivoGen) was transfected with ViaFect™ Transfection Reagent as described above. As TBK-1 inhibitor BX-795, (S1274, Selleck Chemicals) was used. BX-795 stock was diluted in DMSO and a stock concentration of 10mM was generated. CCCP powder (C2759, Sigma-Aldrich) was diluted in DMSO to generate a stock concentration of 100 mM. The protease inhibitors pepstatin A1 (ALX-260-085, Enzo Life Science) and E64d (BML-PI107, Enzo Life Science) were diluted to generate a 10 mg/mL stock concentration in DMSO and Ethanol, respectively. Nuclei were stained with Hoechst 33342 (Invitrogen, H21492). To measure mitochondrial membrane potential, cells were stained with 5 µM JC-1 dye (T3168, Molecular Probes) for 30 minutes at 37°C, washed twice with D-PBS for 5 min, followed by addition of live cell imaging solution (A14291DJ, Thermo Fisher). The cells were illuminated at 488 nm and the emission was measured between 515/545 nm and 575/625 nm with mithras^2^ (Berthold Technologies GmbH & Co. KG, Bad Wildbad Germany) or the Safire II plate reader (Tecan, Maennedorf, Switzerland). Mock infected- or no agnoprotein-expressing cells were set to 100%.

### Antibodies

Goat anti-mouse IgG2a Alexa Fluor 568 (A21134, Invitrogen), donkey anti-rabbit Alexa Fluor 647 (Abcam, ab150075), goat anti-mouse IgG1 Alexa Fluor 488 (A-21121, Molecular Probe), goat anti-mouse IgG1 Alexa Fluor 647 (A-21240, Invitrogen), goat anti-mouse IgG2b Alexa Fluor 647 (A21242, Molecular Probe), goat anti-chicken Alexa Fluor 488 (A11039, Molecular Probes), goat anti-rabbit Alexa Fluor 488 (ab150077, Abcam), donkey anti-guinea pig IRDye 680RD (926-68077, LI-COR) donkey anti-mouse Alexa Fluor 680 (A10038, Invitrogen), goat anti-rabbit IRDye 800CW (926-32211, LI-COR), donkey anti-goat IRDye 800CW (926-32214, LI-COR), polyclonal rabbit anti-aa40-59 JCPyV agno-sera (generated on request by Eurogentec, Belgium), polyconal anti-BKPyV agnosera (generous gift from C. Hanssen-Rinaldo, clone 81038), polyclonal rabbit anti-aa52-66 BKPyV agno-sera (generated on request by Eurogentec, Belgium, clone 753), polyclonal rabbit anti-aa40-53 BKPyV agno-sera (generated on request by Eurogentec, clone 1163), polyclonal rabbit anti-BKPyV LTag sera (generous gift from C. Hanssen-Rinaldo, clone81048), mouse monoclonal IgG2a anti-SV40-LTag cross-reacting with BKPyV LTag (PAb416, Calbiochem), mouse monoclonal IgG1 anti-BKPyV Vp1 (MAB3204-M19, Abnova), polyclonal rabbit anti-SV40 Vp2/Vp3 (ab53983, Abcam), mouse monoclonal IgG2a anti-Tom20 (sc-17764, Santa Cruz), mouse monoclonal IgG2a anti-MAVS (sc-166583, Santa Cruz), mouse monoclonal IgG1 anti-IRF3 (NBP1-04308, Novus Biologicals), mouse monoclonal IgG1 anti-Interferon *α* (21116-1, pbl assay science), mouse monoclonal IgG1 anti-Interferon *β* (21400-1, pbl assay science), mouse monoclonal IgG2a anti-Interferon *α/β* receptor chain 2 (21385-1, pbl assay science), mouse monoclonal IgG1 anti-p62/SQSTM1 (sc-28359, Santa Cruz), polyclonal guinea pig anti-p62/SQSTM1 (GP62-C, Progen), mouse monoclonal IgG1 anti-PDI (SPA-891, Enzo Life Science), polyclonal chicken anti-Calreticulin (ab14234, Abcam), polyclonal rabbit anti-Phospho-Drp1-(Ser616) (3455, Cell Signalling), mouse monoclonal IgG1 anti-Drp1 (NBP2-23489, Novus Biologicals), mouse monoclonal IgG2b anti-STING (MAB7169, R&D Systems), polyclonal rabbit anti-LC3A/B (4108S, Cell Signalling), mouse monoclonal IgG1 anti-beta Actin (ab6276, Abcam), mouse monoclonal IgG1 and anti-Tubulin (A-11126, Molecular Probes).

### Cell lines and viruses

Primary human renal proximal tubular epithelial cells (RPTEC Lot:5111; 4100, ScienCell) were maintained in epithelial cell medium (EpiCM; 4101, ScienCell) and passaged with Trypsin (T3924, Sigma-Aldrich) and Defined Trypsin Inhibitor (DTI; R007100, Invitrogen). UTA6-2C9 cells stably transfected with the *AGN* gene under a tetracyclin-dependent suppressor (tet-off) have been described previously (Cioni et al., 2013).The African green monkey kidney cell line CV-1 (ATCC CCL-70) was cultured in Dulbecco’s modified Eagle’s medium (D5671, Sigma-Aldrich) supplemented with 5% FBS (S0115, Biochrom) and 2mM Stable Glutamine (5-10K50-H, Amimed). The SVG-A cells (generous gift from C. Hanssen-Rinaldo) were kept in Minimum Essential Medium (M2279, Sigma-Aldrich) including 10% FBS. One day after seeding, RPTEC were infected with BKPyV Dunlop at a MOI of 1.0 determined by nuclear LTag staining on RPTECs. The infection was carried out for 2 h before surplus infectious units were removed and EpiCM containing 0.5% FCS was added. One day after seeding, CV-1 cells were infected with SV40 (776 strain) at a MOI of 0.5-1.0 pfu/cell using a supernatant from SV40-infected BS-C-1 cells. The infection was carried out for 2 h before surplus infectious units were removed and complete medium added. One day after seeding, SVG-A cells were infected with JCPyV Mad-4 using a supernatant from JCPyV Mad-4 transfected COS-7 cells. The infection was carried out for 2 h before surplus infectious units were removed and complete medium added.

### Plasmids

The different agnoprotein constructs have been described in (Unterstab et al., 2010). Mitochondrial membrane localization signal (MTS) of cytochrome c oxidase cox8 was N-terminally fused to agno(20-66)-EGFP and named MTScox8-Agno(20-66). The mCherry-mEGFP-OMP25TM tandem tag mitophagy reporter was a generous gift of Professor Terje Johansen, UiT The Arctic University of Norway, Tromsø, Norway. The YFP-Parkin plasmid was a gift from Richard Youle (Addgene plasmid # 23955; http://n2t.net/addgene:23955; RRID:Addgene_23955)(Narendra et al., 2008).

### Electron microscopy

Samples were rinsed with ice-cold cacodylate 0.1M pH7.4 and fixed with 2% paraformaldehyde, 2.5% glutaraldehyde (01909; Polysciences) in 0.1M cacodylate pH7.4. Samples were washed again with cacodylate buffer (0.1M, pH=7.0 and postfixed in 1% osmium tetroxide and 1.5% potassium ferrocyanide in cacodylate buffer (0.1 M pH=7.4), followed by a 1% osmium in cacolylate buffer treatment. Sections were washed in distilled water. Samples were stained with 1% uracil acetate in water and rinsed once with H_2_O. Samples were de-hydrated in graded alcohol series and embedded in Epon. The images were taken with a TecnaiTm Spirit TEM (Fei, Thermo Fisher). Biopsy specimens were processed and analyzed as described previously (Drachenberg et al., 1999).

### Immunofluorescence microscopy

Microscopy was performed using an epifluorescence microscope (model eclipseE800; Nikon, Tokio, Japan) equipped with suitable filters and a digital camera (Hamamatsu, Tokio, Japan). The pictures were analyzed by Fiji (U.S. National Institutes of Health, Bethesda, MD) as described previously (Hirsch et al., 2016).

### Confocal Laser Scanning Microscopy (CLSM)

Confocal pictures were taken with a LeicaSP5 (Leica, Wetzlar, Germany) with a 63x Plan Apochromat/NA1.4 oil objective and pictures were further processed by Fiji. Z-stacks were acquired with a 63x Plan Apochromat/NA1.4 oil objective and the requirements of the Nyquist theorem were fulfilled, with voxel *xyz* size of 45, 45, 150 nm, respectively. To exclude the possibility of channel crosstalk, images were acquired sequentially using the multi-track mode. Deconvolution and visualization were done essentially as described in (Unterstab et al., 2010). For deconvolution, the Huygens professional software (Scientific Volume Imaging, Hilversum, the Netherlands) was used, applying the Classic Maximum Likelihood Estimation Mode (CMLE) using a theoretical point spread function (PSF) and IMARIS (Bitplane AG, Zurich, Switzerland) was used for visualization, colocalization analyzes, and quantification.

### Transfection

Transfection of BKPyV genomic DNA or plasmids containing the agnoprotein into RPTECs or UTA6 cells was performed at 90-95% confluency using ViaFect™ Transfection Reagent (E4982, Promega) at a reagent: DNA ratio of 3:1 according to manufacturer’s instructions. At 24 h post-transfection, medium was replaced with EpiCM and DMEM high Glucose containing 5% FBS (S0115, Biochrom AG), respectively. For p62/SQSTM1 knock-down experiments the SMARTpool siGENOME SQSTM1 siRNA (M-010230-00-0005, Dharmacon) and siGENOME Non-Targeting siRNA #2 (D-001210-02-05, Dharmacon) were transfected with Lipofectamine 3000 Reagent (L3000001, Thermo Fisher) according to manufacturer’s instructions, using 100 nM siRNA.

### SDS/PAGE and immunoblotting

Cells were lysed with RIPA buffer: 10 mM Tris/HCl pH 7.5; 150mM NaCl; 0.5mM EDTA. 1.0% Nonidet P-40 and proteinase inhibitor (04693132001, Roche). Cell lysates were separated by Mini Protean TGX Gradient Gel 4-20% (4561095, Bio-Rad) and electrotransferred onto 0.45-μ Immobilon-FL polyvinylidene difluoride (PVDF) membrane (IPFL00010, Millipore/Merck). Membranes were blocked with Odyssey blocking buffer (927-40000, LI-COR) diluted 1:2 in Tris-buffered saline (TBS). Primary and secondary antibodies were diluted in Odyssey blocking buffer diluted 1:2 in TBS-0.1% Tween 20 and incubated at RT for 1 h (or o/n at 4°C) or 45 min, respectively. Washing in between was performed with TBS–0.1% Tween 20. For the detection of p62/SQSTM1, cells were trypsinized and 2.0×10^4^ cells were resuspended and lysed per 10µL 1x Laemmli Sample Buffer (161-0747, Bio-Rad). 10µL cell lysate was separated by SDS-PAGE, electrotransfered onto 0.2-µm Immobilon®-P^SQ^ PVDF membrane (ISEQ00010, Millipore/Merck) and immunoblotting was done essentially as described above, but 3.0% milk (T145.2, Roth) in TBS-0.1% Tween 20 was used instead of the Odyseey blocking buffer. Detection and quantification were done with the Odyssey CLx system (LI-COR, Lincoln, USA).

### Enzyme-linked immunosorbent assays (ELISA)

The ELISA was performed using the VeriKine-HsTM Human IFN Beta Serum ELISA KIT (41415, pbl assay science). The recommended enhanced protocol form improved performance in serum evolution was used. Samples were measured by Safire II plate reader in triplicates and the standard curve was generated using GraphPad Prism software (v8.1.0).

### Immunohistochemistry

After deparaffinization samples were heated for 30min in the microwave at 98°C in citrate-buffer (pH 6.0) and cooled down for 30min. Samples were washed 1*×*5min with D-PBS and 1*×*5min in 10mM D-PBS + 0.1% Tween20 (93773, Fluka). Samples were blocked with 5% normal goat serum (50197Z, Thermo Fisher), for 1h at RT. Primary antibodies were diluted in 3% BSA/PBS (A9647, Sigma-Aldrich) and samples were incubated over night at 4°C followed by 2*×*5min washes in 10mM PBS. Secondary antibodies were diluted in 3% BSA/PBS containing 1µg/ml Hoechst 33342 (B2261, Sigma-Aldrich), followed by 2*×*5min washes in 10mM PBS. Samples were mounted with Prolong Gold (P36935, Thermo Fisher).

### Quantitative nucleic acid testing (QNAT)

BKPyV loads were quantified form cell culture supernatant as described previously (Hirsch et al., 2016). To quantify interferon-*β* transcripts, cellular RNA was extracted using the QIAshredder kit and the RNeasy Mini kit (79654 and 74106, Qiagen). The cDNA was generated using High Capacity cDNA reverse transcript kit (4368814, Applied Biosystems) and RT-QNAT was performed on a Veriti 96 Well Thermal Cycler (Applied Biosystems, Lincoln, USA) using an human interferon b1 assay (4331182, Thermo Fisher Scientific) and human HPRT1 endogenous control (4333768F, Applied Biosystems).

### Statistics

All results were analyzed by GraphPad Prism software (version 8.1.0). Level of significant variability between groups was tested as indicated in figure legends, using paired t-test or two-way analyses of variance (ANOVA), respectively. Significant differences were assumed for P values of <0.05. Unless indicated differently, groups were plotted as mean values and standard deviation (SD).

## Supporting information

Suppl Mov 1

Suppl Mov 2

Suppl Mov 3

## Acknowledgments

We thank Pascal Lorentz of the confocal imaging core facility of the Department Biomedicine, University of Basel, and Mohamed Chami at the electron microscopy core facility of the Biocenter, University of Basel for advice and support with the imaging studies. We are grateful to Professor Terje Johansen, UiT The Arctic University of Norway, Tromsø, Norway, for helpful discussions and for providing the tandem tag mitophagy reporter construct mCherry-mEGFP-OMP25TM. We thank Ms Erika Hofmann for the timely updates of the reference library. The data presented in this manuscript are tabulated in the main paper and in the supplementary materials.

## Funding

This study was funded by an appointment grant of the University of Basel to H.H.H.

## Author contributions

J Manzetti, G Unterstab, F Weissbach, M Wernli, C Hanssen-Rinaldo, C Drachenberg, H Hopfer conducted experiments, analyzed data, and contributed to writing the manuscript; HHH designed the study, analyzed data, and wrote the manuscript.

## Declaration of Interests

The authors declare no competing interests

**Supplementary Material 1: Figure S1.**

**BKPyV protein expression and intracellular colocalization in primary human RPTECs** Cells were infected with the indicated viral strains, fixed at 48 hpi, stained for confocal microscopy and z-stack acquisition as described in Materials & Methods. (a) Dun-*AGN* LTag. (red), Vp1 (cyan), agnoprotein (green), and DNA (blue). (b) Dun-*agn25D39E.* LTag (red), Vp1 (cyan), agnoprotein (green), and DNA (blue). (c) Dun-*agn25D39E*. Tom20 (red), agnoprotein(green), PDI (magenta) and DNA (blue). Colocalizing voxels are shown in yellow.

**Supplementary Material 2: Figure S2.**

**BKPyV Dun-*AGN*-infected RPTECs showTom20-p62/SQSTM1 colocalization which is not observed in BKPyV Dun-*agn25D39E* infected RPTECs** RPTECs were infected with BKPyV Dun-*AGN* or Dun-*agn25D39E* and at 72 hpi confocal microscopy was performed for Tom20 (red) agnoprotein (green), p62/SQSTM1 (cyan), and DNA (blue). Colocalizing voxels shown in yellow.

**Supplementary Material 3: Figure S3.**

**UTA6-2C9 cells transfected with YFP-Parkin show parkin-positive mitophagy following CCCP treatment, but not following agnoprotein expression.** UTA6-2C9 cells harboring the tetracycline (tet)-off inducible BKPyV agnoprotein were transfected with yellow-fluorescence protein-Parkin construct, and confocal microscopy was performed at 48 hpt.

BKPyV agnoprotein suppression (tet+) and expression (tet-) is indicated, respectively. CCCP treatment (10µM) for 3 h served as positive control for parkin-positive mitophagy. Confocal images of cells stained for DNA (blue), Tom20 (red), YFP-parkin (yellow), and agnoprotein (green). Magnification and 3D-isosurface rendering.

**Supplementary Material 4: Figure S4.**

**JCPyV agnoprotein colocalizes to mitochondria and induces network fragmentation.**
Z-stacks of JCPyV infected SVG-A cells were acquired, deconvolved and processed with IMARIS. 3D isosurface-renderings of the mitochondrial marker Tom20 (red), agnoprotein (green), and DNA (blue) are shown. Colocalizing voxels are shown in yellow.

**Supplementary Material 5: Video S1. Dun-AGN Agnoprotein colocalizes with mitochondria and induces mitochondrial fragmentation.** Cells were infected with the indicated viral strains and fixed at 48 hpi as described in Materials & Methods. Z-stacks of BKPyV Dun-*AGN*, stained for Tom20 (red), agnoprotein (green), and DNA (blue). Colocalising voxels are shown in yellow.

**Supplementary Material 6: Video S2. Dun-*agn25D39E* mutant agnoprotein colocalizes with mitochondria, but does not induces fragmentation of mitochondrial network.** Z-stacks of BKPyV Dun-*agn25D39E* infected cells, stained for mitochondrial marker Tom20 (red), agnoprotein (green), and DNA (blue). Colocalizing voxels are shown in yellow.

**Supplementary Material 7: Video S3. MTScox8-agno-(20-66)-mEGFP is targeted to mitochondria and induced mitochondrial fragmentation.** UTA6 cells transfected with MTScox8-agno(20-66) at 24 hpt. Immunofluorescent staining for Tom20 (red) and agnoprotein-mEGFP (green); DNA (blue). Colocalizing voxels are shown in yellow.

## Notes

#### Summary of Updates

This paper correspond to the non-peer reviewed version.

