## Supplementary figures and images for "BK polyomavirus evades innate immune sensing by disrupting the mitochondrial network and membrane potential and by promoting mitophagy"

### Suppl Fig S3

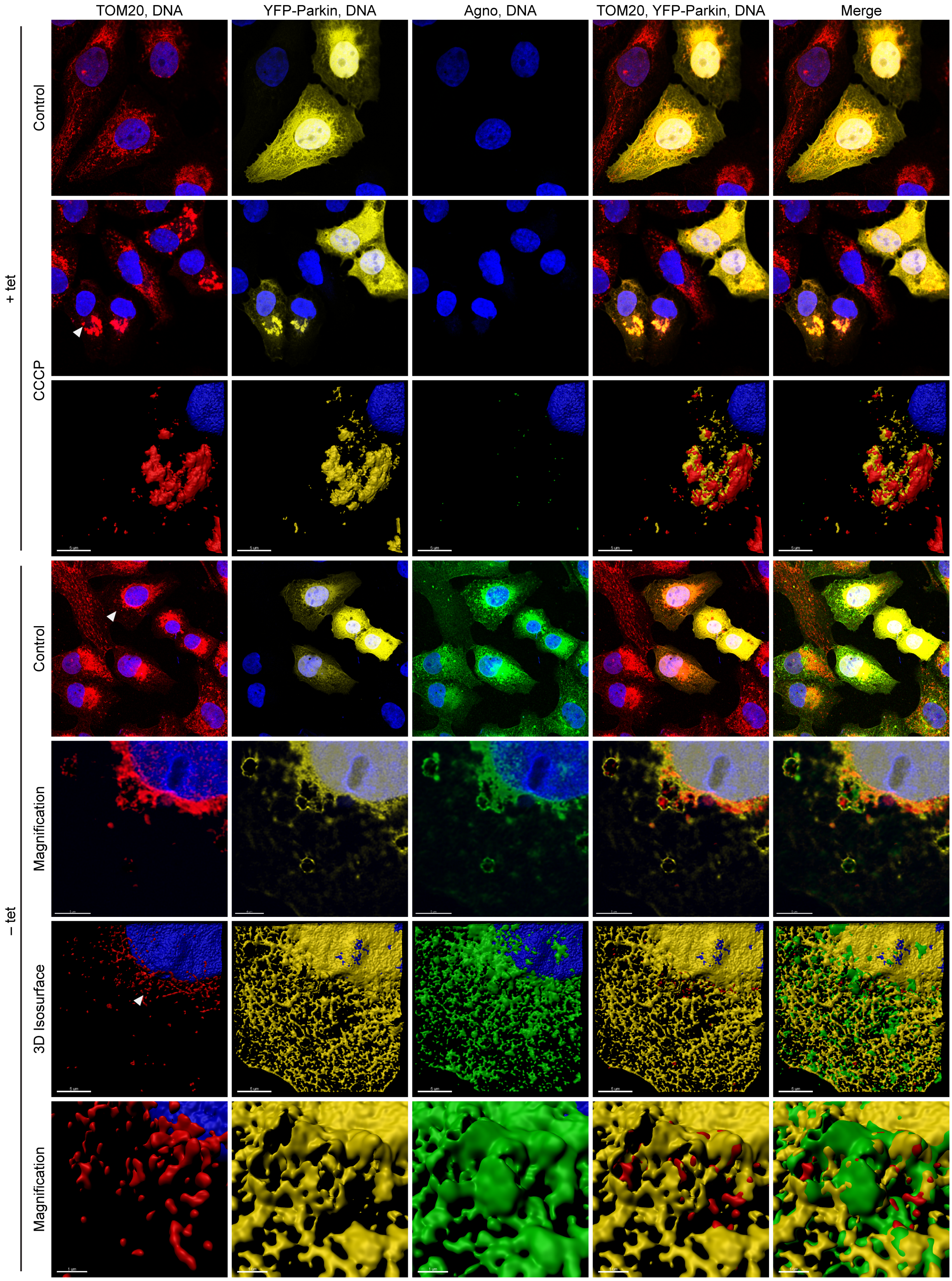

### Suppl Fig S4

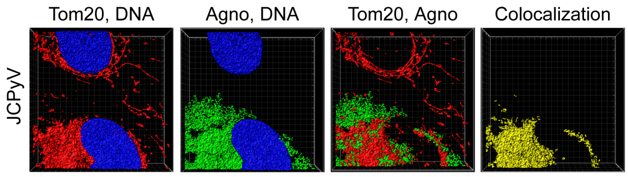

### Suppl FigS1

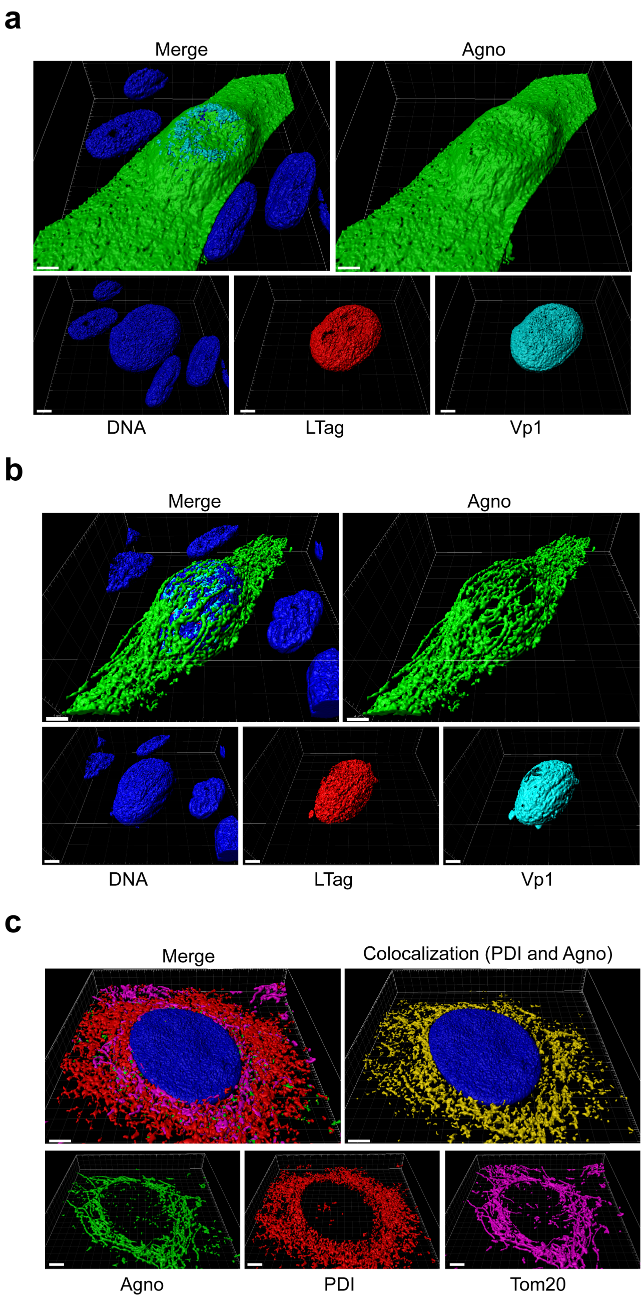

### Suppl FigureS6.tif

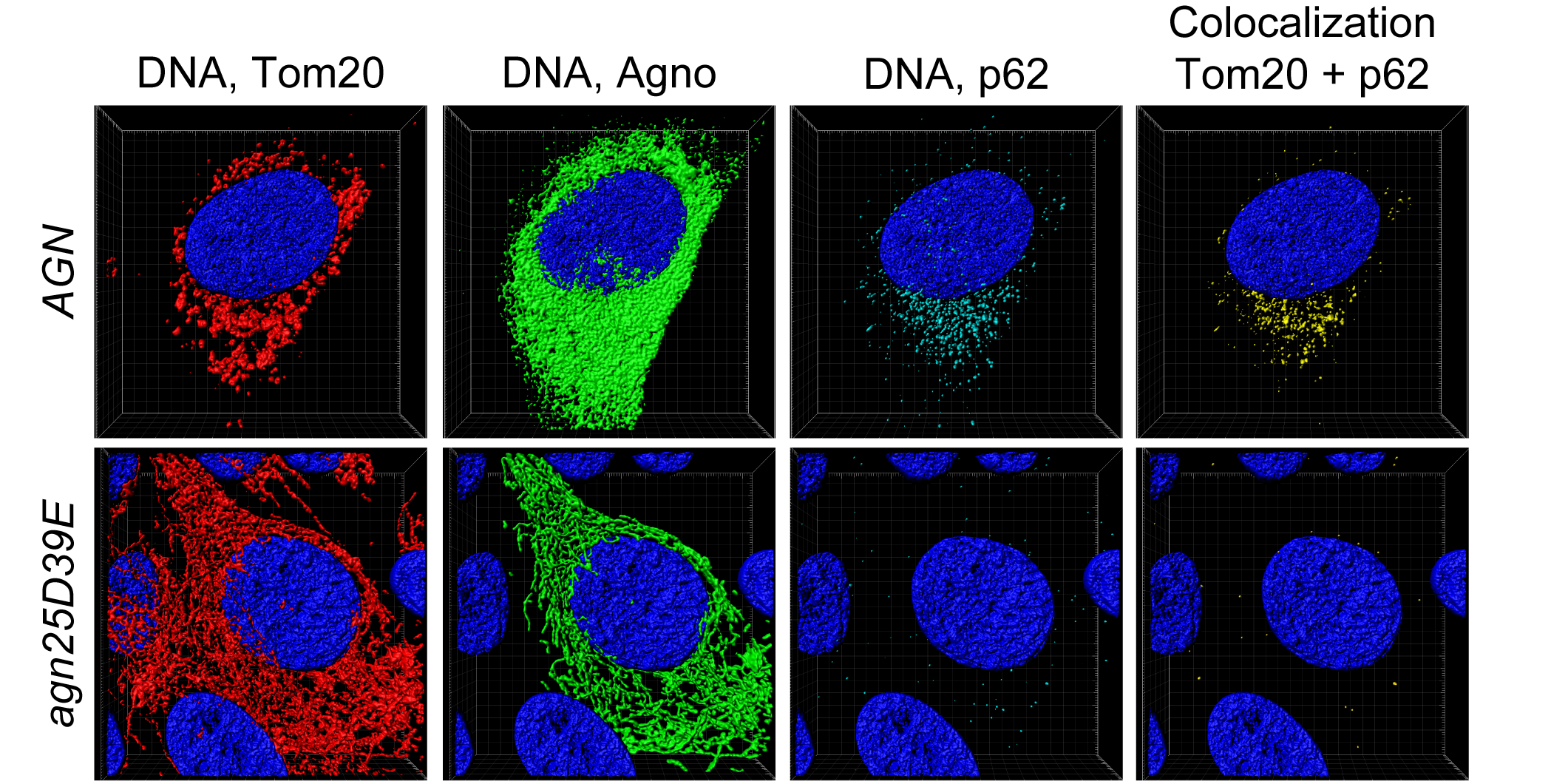
